# Cryo-electron tomography of NLRP3-activated ASC complexes reveals organelle co-localization

**DOI:** 10.1101/2021.09.20.461078

**Authors:** Yangci Liu, Haoming Zhai, Helen Alemayehu, Jérôme Boulanger, Lee J. Hopkins, Alicia C. Borgeaud, Christina Heroven, Jonathan D. Howe, Kendra E. Leigh, Clare E. Bryant, Yorgo Modis

## Abstract

NLRP3 induces caspase-1-dependent pyroptotic cell death to drive inflammation. Aberrant activity of NLRP3 occurs in many human diseases. NLRP3 activation induces ASC polymerization into a single, micron-scale perinuclear punctum. Higher resolution imaging of this signaling platform is needed to understand how it induces pyroptosis. Here, we apply correlative cryo-light microscopy and cryo-electron tomography to visualize ASC/caspase-1 in NLRP3-activated cells. The puncta are composed of branched ASC filaments, with a tubular core formed by the pyrin domain. Ribosomes and Golgi-like vesicles permeate the filament network, consistent with roles for these organelles in NLRP3 activation. Mitochondria are not associated with ASC but have outer-membrane discontinuities the same size as gasdermin D pores, consistent with our data showing gasdermin D associates with mitochondria and contributes to mitochondrial depolarization.

**One-Sentence Summary:** Electron tomography of frozen cells reveals the ultrastructure of ASC specks and gasdermin D pores in adjacent mitochondria.

## Main Text

Inflammasomes are oligomeric signaling complexes that induce proteolytic activation of caspase-1, leading to processing of the proinflammatory cytokines interleukin (IL)- 1β and IL-18, and cleavage of the cell death effector gasdermin D (GsdmD) to induce pyroptosis. In the first step of inflammasome assembly, Nod-like receptors (NLR) sense cytosolic chemical signatures associated with microbes and cell dysfunction or damage (*1*). Ligand binding induces NLR oligomerization and recruitment of downstream proteins linked to the caspase-1 signaling cascade.

NLRP3 is one of the most studied inflammasome NLRs due to its association with many human diseases including degenerative disorders, dermatitis, and metabolic disorders (*2, 3*). Over 250 human NLRP3 protein variants have been identified, of which at least 30 have been associated with autoinflammatory disease through gain-of-function (*4, 5*). NLRP3 inflammasome activation has been linked to a broad spectrum of cellular processes or cytosolic chemical signatures including potassium efflux, reactive oxygen species, ribosomal arrest, calcium influx, chloride efflux, phosphatidylinositol-4-phosphate on Golgi membranes, and mitochondrial stress or damage (*6, 7*). In addition to ligand recognition, activation of the NLRP3 and pyrin inflammasomes may require active transport along microtubules by dynein to the microtubule organizing center (MTOC) (*8, 9*). Precisely how these different activation mechanisms and the associated involvement of subcellular organelles lead to NLRP3 activation remains to be elucidated.

Cryo-electron microscopy (cryo-EM) image reconstructions show that ligand-bound inflammasome NLRs assemble into disk- or split washer-like oligomers (*10-13*). Upon activation, NLRP3 recruits the adaptor protein ASC (Apoptosis-associated speck-like protein containing a caspase recruitment domain (CARD)). NLRP3 and ASC interact through their pyrin domains (PYDs) (*14*). ASC then can assemble into a single perinuclear punctum with a micron-scale diameter, also known as a speck, which functions as a platform for the recruitment and activation of caspase-1 (*15, 16*). The PYD and CARD of ASC are both required for punctum formation and signaling (*15, 17, 18*). Structural studies on purified ASC PYD and ASC CARD show that both domains have a death fold and independently form helical filaments (*19-24*). In the cell, fluorescence microscopy imaging suggests that PYD-PYD interactions drive ASC filament formation whereas CARD-CARD interactions promote cross-linking and compaction of PYD filaments into puncta (*17, 18, 23-26*). In addition, ASC recruits procaspase-1 via a CARD-CARD interaction between the two proteins (*15, 23*). Super-resolution light microscopy of endogenous ASC specks revealed punctate structures containing ASC, caspase 1 and NLRP3 within the same macro-assembly (*27*). These structures have yet to be imaged inside a cell at sufficient resolution to resolve the ASC or caspase-1 ultrastructure within puncta (*26*). Hence, it remains unclear how the PYD and CARD contribute to the formation of an ASC punctum inside cells.

Understanding how ASC filament formation contributes to the assembly and activation of inflammasome puncta requires structural information in the cellular context. Recent advances in the preparation of frozen vitrified cellular lamellae by focused ion-beam (FIB) milling (*28, 29*) have allowed the ultrastructure of multimeric protein complexes to be determined in a near-native context by cryo-electron tomography (cryo-ET). Here, we perform *in situ* cryo-ET to obtain three-dimensional image reconstructions of multimeric signaling complexes in cells with active NLRP3 inflammasomes. We identify ASC/caspase-1 signalosome puncta by correlative light and electron microscopy (CLEM) and reveal their ultrastructure and pyroptotic cellular landscape in immortalized bone marrow derived macrophages (iBMDMs) by cryo-ET image reconstruction. We find that inflammasome puncta are composed of short, hollow ASC/caspase-1 filaments, some branched, with variable packing density allowing ribosomes and *trans-*Golgi-lie vesicles to permeate the puncta. Neither the MTOC nor mitochondria are associated with the ASC speck although the ultrastructural analysis indicates local disruption or pore formation in the outer mitochondrial membrane. Time resolved analysis of NLRP3 stimulated ASC speck formation shows it occurs concurrently with loss of mitochondrial integrity. We propose that the ultrastructure of the ASC filament network functions as a signaling platform by providing structural integrity while allowing downstream signaling molecules to diffuse freely and bind at high density within the network.

## Results

### In situ cryo-CLEM of ASC/caspase-1 puncta

High-resolution structures of inflammasome components have been obtained from purified proteins. In a previous study in zebrafish larvae, electron tomography of ASC puncta in fixed larval sections stained with uranyl acetate showed that the puncta contained a dense filament network (*26*), but the imaging resolution was limited by chemical fixation and staining of the sample. To examine the architecture and cellular interactions of the NLRP3 signalosome in its physiological environment, we imaged ASC puncta in iBMDMs by correlative fluorescence light microscopy and *in situ* cryo-ET (cryo-CLEM). Cells expressing ASC fused with a C-terminal fluorescent protein have been used extensively to visualize ASC puncta formation (*30*). In some experimental systems, ASC overexpression can lead to puncta formation in the absence of activating inflammasome stimuli (*30*). Fluorescence microscopy of live iBMDM cells overexpressing ASC-mCerulean used in this study showed that both priming with lipopolysaccharide (LPS) and stimulation with nigericin were still required to induce ASC puncta formation (**Fig. S1A**). Time course analysis showed ASC speck formation occurring within 20 min of nigericin stimulation (**Movie S1**). GsdmD cleavage observed by Western blotting and IL-1β cytokine release measured in ELISAs confirmed that the ASC-mCerulean puncta activated pyroptotic signaling (**Fig. S1B**). Limiting the stimulation time with nigericin to 30 min allowed early-stage inflammasome signaling complexes to be captured, as indicated by colocalization of NLRP3, proteolytically activated caspase-1 and IL-1β with ASC in the puncta, in both wild-type (WT) iBMDMs and ASC-mCerulean iBMDMs (**Fig. S1C-E**). NLRP3 and pro-IL-1β localization was determined by immunofluorescence. Caspase-1 localization was inferred using the carboxyfluorescein-labeled caspase-1 substrate FAM-FLICA. FAM-FLICA or its non-fluorescent analogue Z-VAD-FMK also facilitated imaging by delaying pyroptotic cell death (**Movie S1**).

For high-resolution imaging, cells grown on electron microscopy grids were vitrified by plunge-freezing in liquid ethane. We used both iBMDMs overexpressing ASC-mCerulean and WT iBMDMs labeled with FAM-FLICA after NLRP3 inflammasome activation (*31*). ASC/caspase-1 signaling complexes were located by cryo-fluorescence light microscopy (cryo-FM; **Fig. S2A**). Areas of the frozen cells containing an ASC or caspase punctum were FIB-milled down to lamellae 150-300 nm-thick and 15-20 µm wide (*28, 29*), with micro-expansion joints to reduce bending (*32*) (**Fig. S2, B** and **C**). The lamellae were imaged by cryo-CLEM (*33*), using cryo-FM to locate any remaining fluorescent signal from ASC-mCerulean or FAM-FLICA in the lamellae after milling (**Fig. 1, A-C; Fig. S2, D** and **E**). In most cases the ASC punctum was milled away during the milling procedure but about 5% of lamellae retained fluorescent signal. The micron-scale dimensions of the fluorescent puncta allowed cryo-ET tilt-series acquisition at regions of interest with the necessary targeting precision after correlation of the cryo-FM images with low-magnification cryo-EM maps (**Fig. S2, F** and **G**) (*34, 35*). To obtain an internal quality metric for the resulting tomograms, we generated a subtomogram averaging reconstruction of 1,058 ribosomes extracted from five ASC speck tomograms. This yielded a ribosome structure with a resolution of 23 Å and the expected structural features (**Fig. 1, E** and **F; Fig. S2H-J**).

**Fig. 1.**
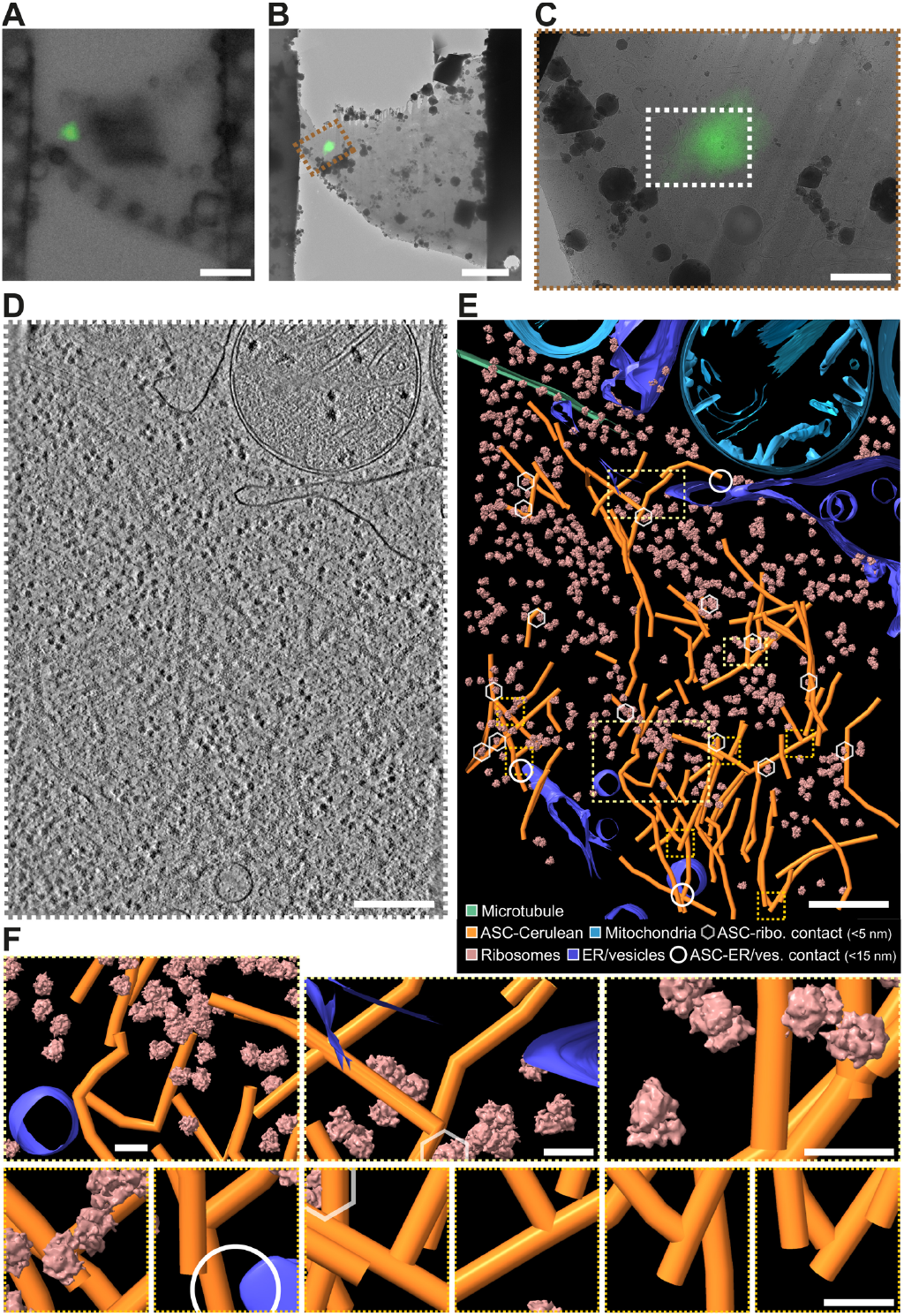
Cryo-ET image reconstructions and models of ASC-caspase 1 puncta in iBMDMs. (**A**) Cryo-FM image of a lamella from an iBMDM expressing ASC-mCerulean. ASC-mCerulean fluorescence is shown (green), overlaid on a bright-field image. Scale bar, 5 µm. (**B**) Cryo-FM image shown in (A) correlated with and overlaid on a cryo-EM map of the lamella with eC-CLEM ICY (*33*). Scale bar, 5 µm. (**C**) Closeup of the area boxed in red in (B). Scale bar, 1 µm. (**D**) Reconstructed cryo-ET volume (13.6-nm thick virtual tomographic slice) of the area boxed in white in (C). Scale bar, 250 nm. (**E**) 3-D segmented model of an 1,873ξ 1,386ξ 108 nm volume covering the area shown in (D), generated with IMOD (*36*). Scale bar, 250 nm. Hexagons denote ASC-ribosome contacts (< 5nm). Circles denote ASC-ER/vesicle contacts (<15 nm). (**F**) Closeups of the boxed areas in (E): Yellow boxes, representative views at different magnifications. Orange boxes, ASC filament branch points. Scale bars, 40 nm. The ribosome structure shown is a 23-Å resolution subtomogram averaging reconstruction of 1,058 ribosomes from five tomograms of ASC-mCerulean puncta (see **Fig. S2**).

### Composition, ultrastructure, and dynamics of ASC/caspase-1 puncta

Three-dimensional cryo-ET image reconstructions show a dense network of branched filaments in areas identified by cryo-CLEM as containing ASC-mCerulean or caspase-1 labeled with FAM-FLICA (**Fig. 1D; Fig. S2G**). A three-dimensional model of the ultrastructure generated with IMOD (*36*) showed that ASC puncta consist of several core regions of densely packed filaments separated by sparser regions, with branching filaments connecting adjacent densely packed regions (**Fig. 1, E** and **F; Movie S2**). Ribosomes were abundant within the filament network. Small vesicles were present, but organelles including mitochondria and the MTOC were excluded. A typical cryo-ET reconstruction (1,900 × 1,400 × 108 nm) contained 10-20 ribosomes in contact with ASC filaments and 2-4 sites with filaments within 15 nm of ER or vesicle membranes (**Fig. 1, E** and **F**). Filaments in cells expressing ASC-mCerulean had additional density at the filament periphery, and a less distinct outline compared to filaments in cells expressing WT ASC (**Fig. 2A**). This suggests the additional density is primarily attributable to mCerulean, which at 27 kDa is larger than ASC (22 kDa), although other components such as caspase-1 (which was shown by light microscopy to localize to ASC specks), may also contribute to this density. A green fluorescent protein tag was similarly found to decorate cryo-ET reconstructions of amyloid-like poly-Gly-Ala aggregates with additional densities (*37*). Since mCerulean was fused to the C-terminus of ASC, the presence of mCerulean at the filament periphery supports a model in which the filaments are formed at their core by the N-terminal PYD of ASC, with the CARD located in between the PYD and mCerulean. Indeed, the cryo-EM structure of purified recombinant ASC PYD filaments and the NMR structure of full-length ASC, taken together, suggest that the PYD forms the filament core (9-nm in diameter), with the flexibly-linked CARD increasing the filament diameter to 16-18 nm (*23, 24*). In our cryo-ET reconstructions, filaments containing WT ASC had a diameter of 11-15 nm, which increased to 22-30 nm for filaments containing ASC-mCerulean (**Fig. 2A**). Notably, a hollow tubular core of density 7 nm in diameter (**Fig. 2, B** and **C**, and **Fig. S3**) was visible within the ASC-mCerulean filaments, for which the cryo-ET images were of higher quality. Neural network image restoration with cryoCARE (*38*) enhanced the definition of the tubular filament core. Although fluorescence from ASC-mCerulean and FAM-FLICA-labeled capsase-1 colocalized in iBMDMs (**Fig. S1E**), the resolution of cryo-CLEM was insufficient to determine whether any of the cryo-ET filament densities corresponded to caspase-1 or other components. We note, however, that caspase-1 (30 kDa) and procaspase-1 (45 kDa) are both larger than ASC and would therefore be expected to increase filament diameter significantly, if present with the same stoichiometry as ASC.

**Fig. 2.**
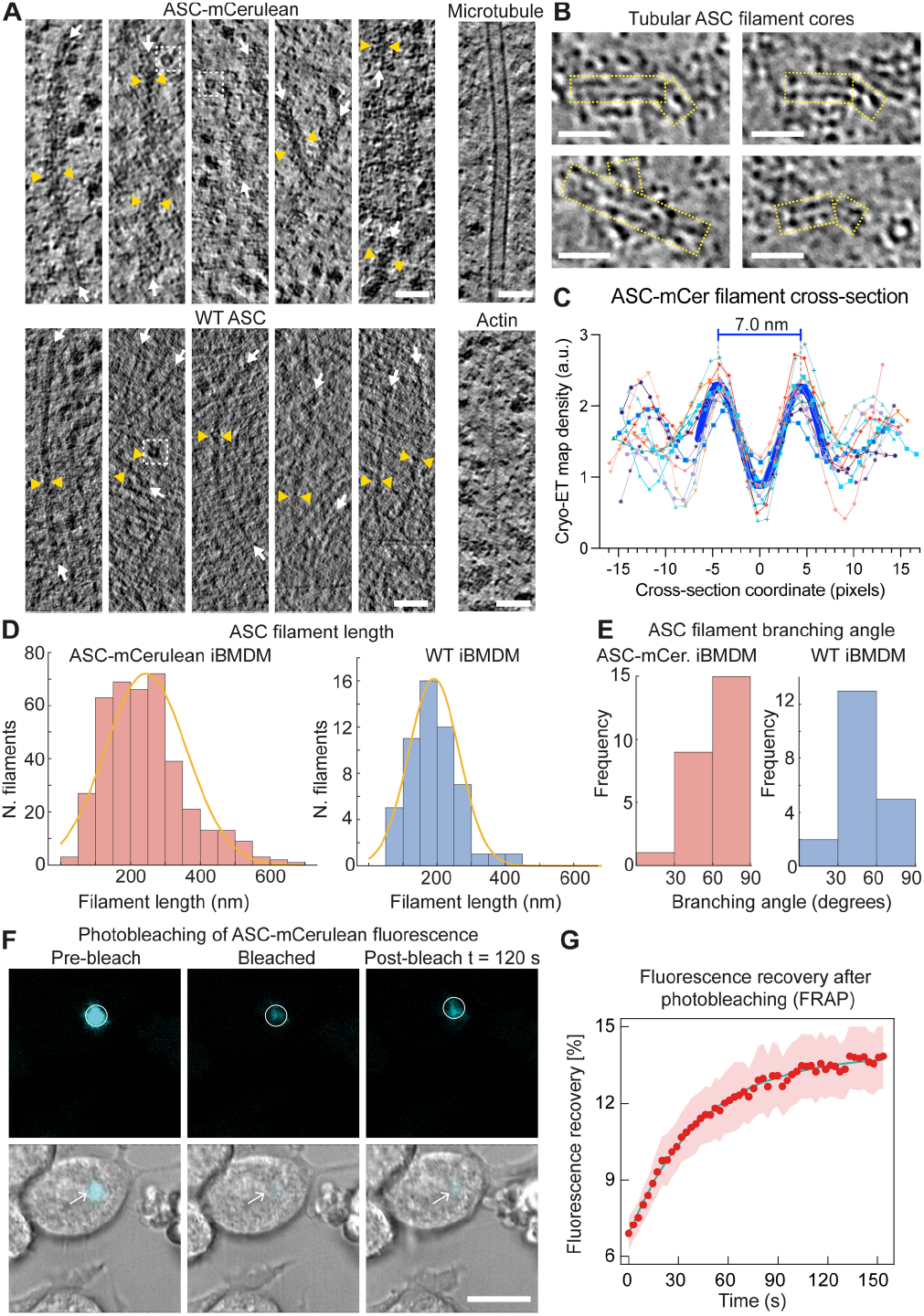
Ultrastructure and dynamics of the ASC filament network in puncta in iBMDMs. (**A**) Filaments in ASC-mCerulean puncta (upper left), or FAM-FLICA-labeled caspase-1 puncta, (lower left), in 4-nm thick virtual tomographic slices. White arrows indicate filament axes. Pairs of yellow triangles show filament thickness. Dashed white boxes highlight ring-shaped densities. Microtubules and actin filaments from the ASC-mCerulean tomogram are shown for reference. Scale bars, 50 nm. (**B**) Tubular filament cores (boxed in yellow) in representative 8-Å thick virtual tomographic slices after image restoration with cryoCARE (*38*). Scale bars, 25 nm. (**C**) Cryo-ET density profiles from 13 ASC filament cross-section areas in 8-Å tomographic slices. 1 pixel = 8 Å. See **Fig. S3** for details on how density profiles were plotted. (**D**) Filament length distribution in ASC-mCerulean iBMDMs (left) and WT iBMDMs labeled with FAM-FLICA (right). (**E**) Distribution of branching angles in ASC-mCerulean puncta (from 2 tomograms), and a WT ASC punctum (1 tomogram). (**F**) Fluorescence recovery after photobleaching (FRAP) of ASC-mCerulean. Scale bar, 10 µm. (**G**) FRAP curve used to calculate the ASC-mCerulean dissociation rate (*k*_off_) from the bleached area. Shaded area represents ± s.e.m. (n = 8). See **Data S1** for source data.

Filament length measurements showed that WT filaments were on average slightly shorter (191 ± 74 nm, n = 54 from one tomogram) than ASC-mCerulean filaments (245 ± 116 nm, n = 402 from six tomograms), with similarly broad length distributions (**Fig. 2D**). This difference could potentially be due to the higher levels of ASC protein expression in iBMDMs expressing ASC-mCerulean, which also express endogenous WT ASC. Analysis of the branching angles showed that the angle distributions of untagged and tagged filaments were slightly different, with larger branching angles more common in mCerulean-tagged filaments, potentially due to steric constraints from the tag (**Fig. 1F, Fig. 2E**, and **Fig. S3**). Our filament branching angle measurements are likely underestimated as any filaments with their axis closely aligned with the Z-axis – which would result in a large (70-90°) branching angle – would not have been detected due to the missing-wedge effect (from sample-tilt limitations). Taken together with previously published structural data from purified proteins, our cryo-ET reconstructions suggest ASC filaments form the backbone of inflammasome-induced puncta, but that the branching of these filaments results in a structural scaffold that can recruit and concentrate downstream effector proteins.

We next investigated the equilibrium dynamics of ASC-mCerulean filaments by fluorescence recovery after photobleaching (FRAP) in iBMDMs. We bleached circular regions containing ASC-mCerulean puncta and imaged the ASC-mCerulean fluorescence intensity before and after bleaching (**Fig. 2F**). Our data showed that ASC-mCerulean filament assemblies have a small mobile fraction (6.9%; **Fig. 2G**). The dissociation rate (*k*_off_) of ASC-mCerulean from the imaged area was measured from the rate of fluorescence recovery as 0.023 ± 0.001 s^-1^ (**Fig. 2, F** and **G; Data S1**). Most of the observed fluorescence recovery was at the periphery of puncta, suggesting that the ASC-mCerulean filaments inside the puncta are largely immobile and are not in equilibrium with ASC monomers or oligomers in the cytosol (**Fig. 2F**).

### Golgi vesicle localization in ASC specks

How NLRP3 is activated by a broad range of different stimuli – including ion fluxes, reactive oxygen species and mitochondrial damage – remains an open question. A common feature of NLRP3 activation by these different stimuli is that the *trans*-Golgi network (TGN) is dispersed, and NLRP3 locally accumulates at foci within the dispersed TGN (*7, 8*). These NLRP3 foci have been proposed to serve as nucleation sites for ASC and caspase-1 recruitment and activation (*7*). Fluorescent ceramide analogs selectively accumulate in the Golgi (*39*). We used the fluorescent BODIPY TR ceramide analog to monitor Golgi localization during ASC speck formation, using either ASC-mCerulean or FAM-FLICA-labeled capsase-1 fluorescence to visualize the specks in live iBMDMs. BODIPY TR ceramide fluorescence was consistently enriched at sites of speck formation (**Fig. 3A; Movie S3**). Immunofluorescence of the TGN marker TGN38 showed that the TGN was dispersed in WT iBMDM and THP-1 cells, and in ASC-mCerulean iBMDMs (**Fig. 3, B** and **C; Fig. S4B**), consistent with previous studies (*7, 8*). Moreover, NLRP3 substantially colocalized with the TGN marker TGN38 in WT iBMDMs primed with LPS, with or without nigericin stimulation (**Fig. 3B**, and **Movie S4**). BODIPY TR ceramide also partially colocalized with TGN38 (**Fig. S4A**). However, there was little direct overlap between ASC-mCerulean and anti-TGN38 fluorescence (**Fig. 3C**). In our tomographic reconstructions, we observed vesicles with diameters of 50 to 300 nm within the ASC filament network (**Fig. 3, D** and **E; S2G; S4C; Movie S2**). We observed similar vesicles within ASC-mCerulean puncta in chemically fixed samples imaged by room temperature electron tomography (**Fig. S4D-F**). The presence of ceramide and absence of TGN38 in the vesicles within ASC puncta suggests these vesicles are derived from the Golgi but not specifically the TGN. Ribosomes were also found embedded throughout the ASC filament network. Ring-shaped densities were occasionally observed within tomograms of ASC puncta, but the resolution of the reconstructions was insufficient to infer the composition of these features (**Fig. 2A**). Together, our cryo-ET and fluorescence microscopy data supports the model that the Golgi apparatus may serve as a platform for ASC recruitment to multiple NLRP3 foci, which nucleate ASC polymerization, with subsequent branching of ASC filaments driving growth into a micron-scale punctum.

**Fig. 3.**
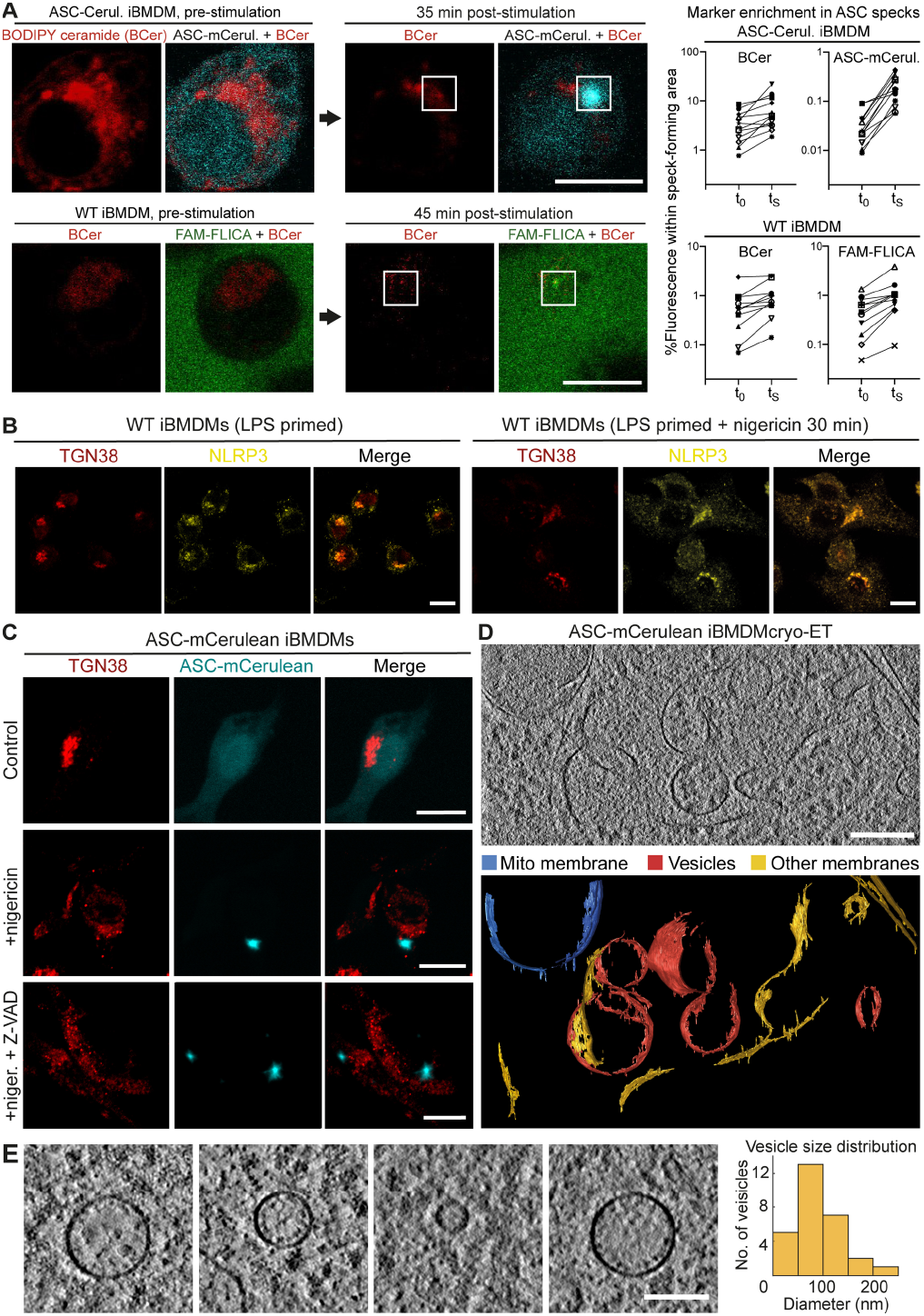
Golgi vesicle localization in ASC specks by live-cell fluorescence confocal microscopy and cryo-ET. (**A**) Distribution of BODIPY TR ceramide (BCer), a Golgi marker, during ASC speck formation in iBMDMs expressing ASC-mCerulean, or WT iBMDMs stained with FAM-FLICA. Areas of speck formation are boxed. See **Movie S3** for full time courses. Right, percentage of whole-cell fluorescence that mapped to the area of speck formation, before and after speck formation, t_0_ and t_S_ (1-3 min and 20-50 min after nigericin addition), respectively. 12 specks from ASC-mCerulean iBMDMs and 11 specks from WT iBMDMs were analyzed. See **Data S2** for source data. (**B**) Immuno-fluorescence microscopy of LPS-primed WT iBMDMs with or without nigericin stimulation. Anti-NLRP3 partially colocalized with anti-TGN38. The TGN was dispersed following stimulation. Scale bar, 10 µm. See **Movie S4** for a Z-stack series. (**C**) Immunofluorescence microscopy of LPS-primed ASC-mCerulean iBMDMs with or without nigericin stimulation, and with nigericin stimulation and caspase inhibitor Z-VAD-FMK. There is little overlap between ASC-mCerulean and anti-TGN38 fluorescence. Scale bar, 10 µm. (**D**) 1.36-nm thick virtual tomographic slice within an ASC speck. Scale bar, 200 nm. Lower panel, 3-D segmentation model. (**E**) Vesicles from cryo-ET reconstructions of ASC-mCerulean iBMDMs. Scale bar, 100 nm. The histogram shows the vesicle size distribution.

Since NLRP3 inflammasome activation was reported to require transport of NLRP3 to the MTOC (*8, 9*), we examined the localization of ASC/caspase-1 puncta relative to the MTOC.

Fluorescent markers for ASC and caspase-1 did not colocalize with the MTOC components γ-tubulin, pericentrin or ninein in WT or ASC-mCerulean iBMDMs stimulated with LPS and nigericin (**Fig. S5**). In stimulated THP-1 cells, ASC/caspase-1 puncta were on average closer to the MTOC, with one quarter of puncta within 2 µm of the MTOC, but there was little direct overlap between the fluorescent markers (**Fig. S5**).

### Aberrant mitochondrial morphology during pyroptosis visualized by cryo-ET

Mitochondria function as a nexus for multiple pathogen-sensing and damage-sensing signaling pathways. Innate immune sensors of viral nucleic acids (*40*) and mitochondrial damage converge on the outer mitochondrial membrane, and induce apoptosis or pyroptosis, depending on the specific danger signal that is sensed (*41, 42*). To visualize and analyze the morphology of mitochondria soon after inflammasome activation we performed cryo-ET reconstructions of mitochondria following LPS priming and nigericin stimulation to induce ASC speck formation. In LPS primed iBMDMs and THP-1 cells expressing WT or ASC-mCerulean ASC, stimulation with nigericin resulted after 30-45 min in mitochondria with a more rounded overall shape and smaller size than in unstimulated cells (**Fig. 4A**, and **Fig. S6**). Cryo-ET reconstructions of iBMDMs and THP-1 cells 1 h post-stimulation showed that the mitochondria in these cells had an abnormal ultrastructure, with the inner membranes forming concentric tubular structures instead of the regularly spaced lamellar cristae observed in unstimulated cells (**Fig. 4A**, and **Fig. S6**). To quantify the changes in mitochondrial morphology associated with ASC-dependent signaling, four parameters were extracted and measured using Fiji (*43*): cristae spacing, inner-to-outer membrane spacing, cristae lumen width and cristae apex angles (**Fig. 4, B** and **C**). We found the cristae spacing to be two- to fourfold smaller in pyroptotic mitochondria than in healthy mitochondria. The other three parameters were not significantly different.

**Fig. 4.**
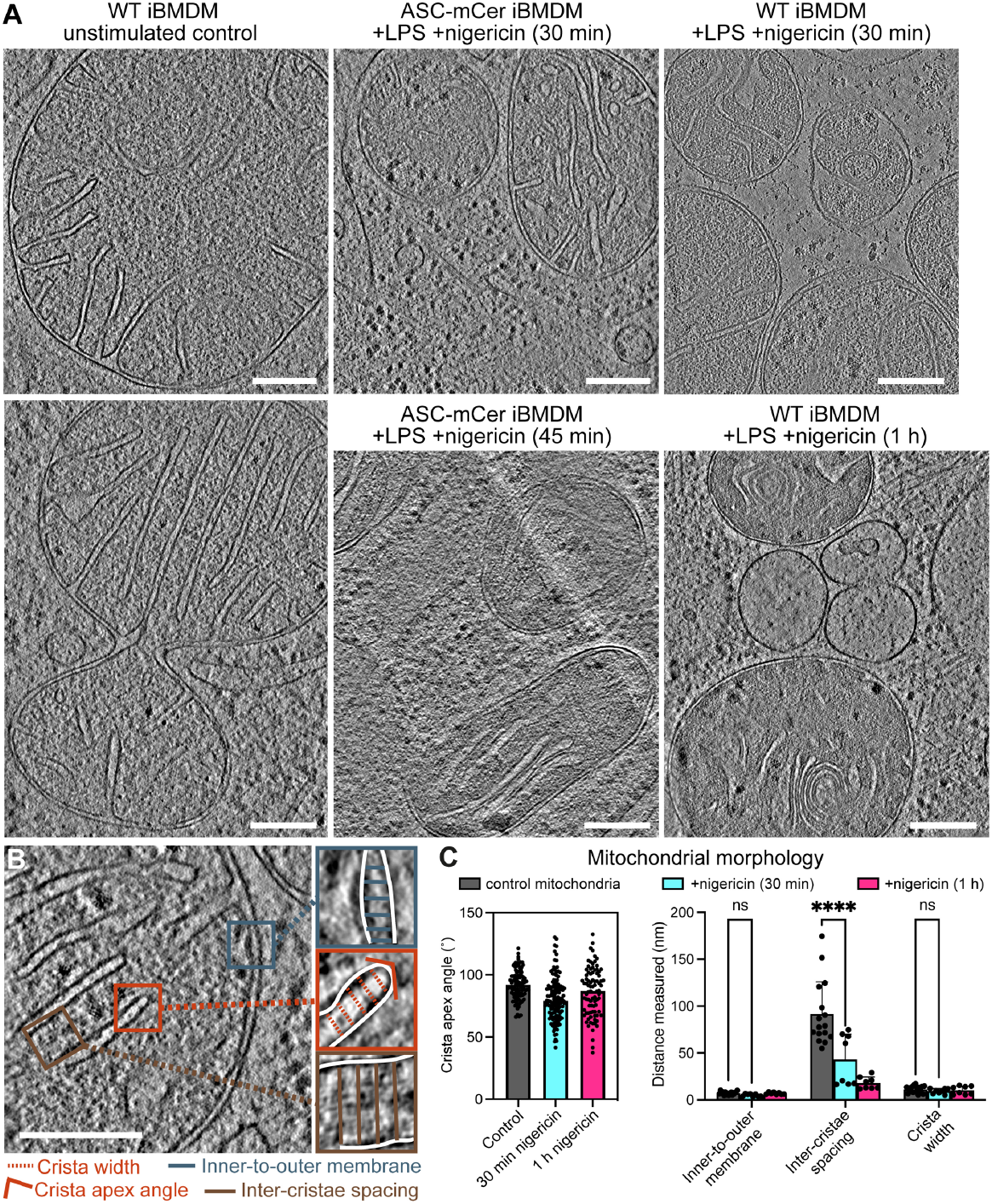
Mitochondrial morphology before and after NLRP3 activation. (**A**) Reconstructed cryo-ET tomographic slices of iBMDMs expressing WT or mCerulean-labeled ASC at different timepoints after stimulation with nigericin. See **Fig. S6** for additional examples including from THP-1 cells. (**B**) Cryo-ET reconstruction of a mitochondrion in an unstimulated WT iBMDM with closeup panels defining the following morphological parameters: inner-to-outer membrane spacing, inter-cristae spacing, crista width and crista apex angle. (**C**) Quantitative analysis of the morphological parameters defined in (B) in segmentation models of the mitochondrial membranes, as shown in (A). Parameters were measured in four tomograms for the control and two tomograms for each of the nigericin-stimulated samples. Error bars represent standard deviation from the mean. For the crista apex angles: “control”, n = 327; “30 min”, n = 187; “1 h”, n = 96. For other parameters: “control”, n = 16; “30 min”, n = 8; “1 h”, n = 8. Statistical test: two-way ANOVA; ns, P > 0.05; ****, P < 10^−4^. Scale bars, 200 nm. See **Data S3** for source data.

### Protein-rich pores 10-20 nm in diameter in the outer mitochondrial membrane

During apoptosis, the Bax and Bak proteins create large disruptions in the outer mitochondrial membrane (*41, 44*). Our cryo-ET reconstructions showed that mitochondria in pyroptotic iBMDMs stimulated with LPS/nigericin lacked any such large outer membrane disruptions. We did, however, identify smaller discontinuities in the outer membranes of some mitochondria in the stimulated ASC-mCerulean and WT iBMDMs (**Fig. 5A**). The size of these gaps, which were only present after nigericin stimulation, varied from 10 to 20 nm (**Fig. 5B** and **Fig. S7A**). The inner membrane was intact near outer membrane gap sites, implying that the gaps were not due to the missing wedge effect or the anisotropic resolution of cryo-ET tomograms (*45*).

**Fig. 5.**
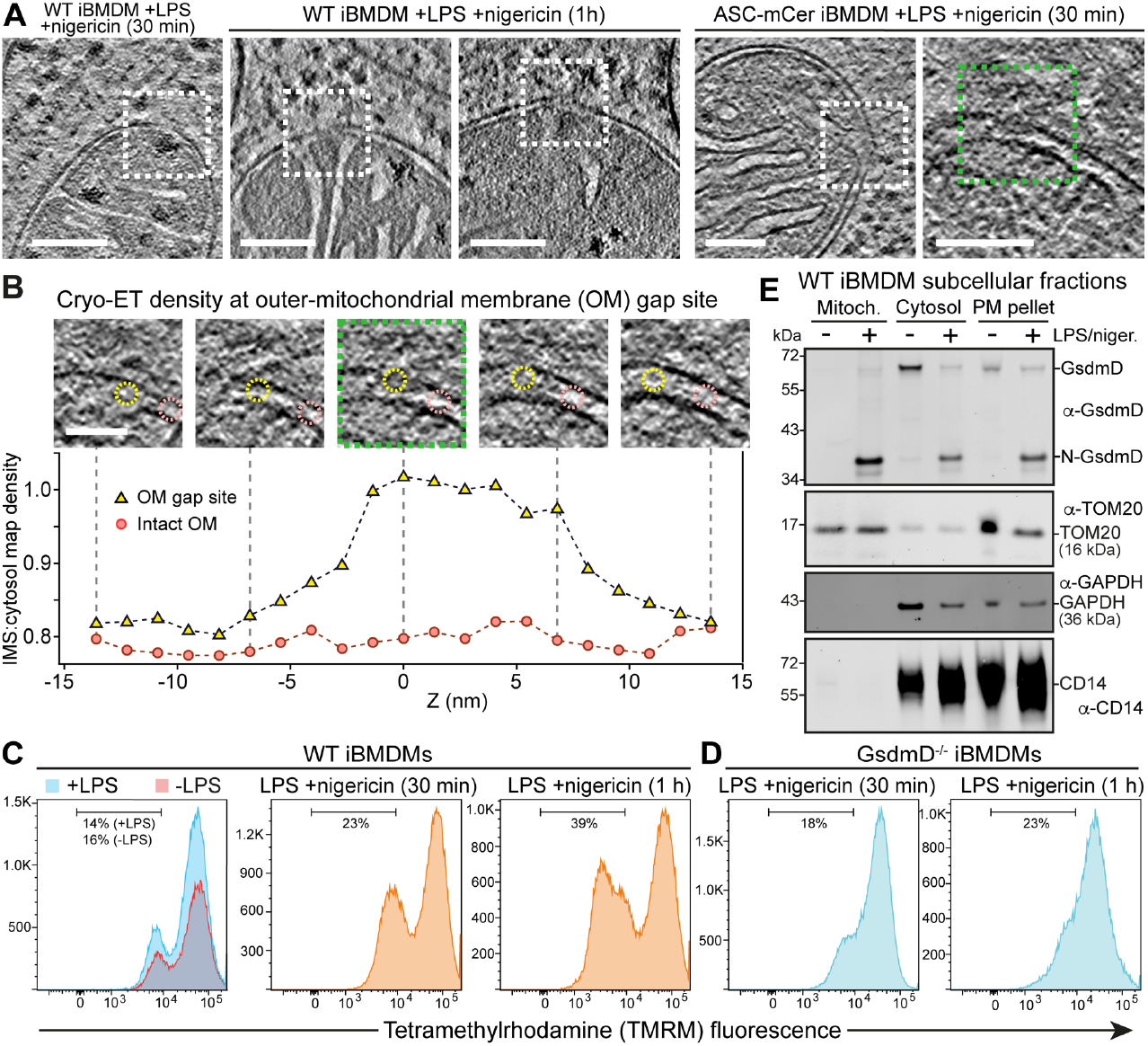
Mitochondrial pore formation, GsdmD-dependent depolarization, GsmdD cleavage and mitochondrial association of N-GsdmD in NLRP3-activated cells. (**A**) Examples of mitochondria with discontinuities in the outer membrane (OM) in cryo-ET reconstructions of in ASC-mCerulean iBMDMs stimulated with LPS and nigericin. Dashed boxes denote the OM discontinuities. Scale bars, 100 nm. (**B**) Z-stack series and cryo-ET density measurements for the OM gap boxed in green in (A). Scale bar, 50 nm. The graph shows the cryo-ET density at the OM gap site (yellow) and at an adjacent site with intact inner and outer membranes (pink). Cryo-ET density is expressed as the density in the intermembrane space (IMS) divided by the density in a nearby cytosolic area. The areas in which IMS density was measured are circled in the Z-stack panels (yellow, gap site; pink, intact site). See **Data S4** for source data. (**C-D**) Flow cytometry histograms of WT iBMDMs, (**C**), or GsdmD^-/-^ iBMDMs, (**D**), stained with tetramethylrhodamine (TMRM), a mitochondrial membrane potential reporter. See **Fig. S8** for additional controls. (**E**) Immunoblots of subcellular fractionation of WT iBMDMs 60 min after stimulation with LPS and nigericin. The GsdmD N-terminal domain (N-GsdmD) is enriched in the mitochondrial fraction. Shown below are immunoblots for Tom20 (a mitochondrial protein), GAPDH (a cytosolic protein), and CD14 (a plasma membrane protein). See also **Fig. S7**.

The cryo-ET map density within outer membrane gaps was lower than that of lipid membranes but higher than in the intermembrane space (**Fig. 5B**). The cryo-ET density at gap sites was greater than in the adjacent intermembrane space over a span of 10-20 nm along the Z-axis, similar to the dimensions of the gaps in X and Y (**Fig. S7A**). This suggests the gap sites contain a higher concentration of protein than the intermembrane space. We conclude that protein-rich gaps or pores, 10-20 nm in diameter, form in the outer mitochondrial membrane during pyroptosis.

### GsdmD pore-forming domain inserts into mitochondria and contributes depolarization

To determine how these outer membrane gaps may contribute to mitochondrial outer membrane permeabilization (MOMP), we quantified membrane potential following inflammasome stimulation in iBMDMs by flow cytometry, using tetramethylrhodamine (TMRM) fluorescence as a reporter. Stimulation with LPS and nigericin or LPS and ATP caused loss of mitochondrial membrane potential in a large fraction of cells (40-50%; **Fig. 5C** and **Fig. S6**).

The N-terminal domains of gasdermins D and E form 20-nm pores in lipid bilayers (*46, 47*), and have been reported to cause mitochondrial depolarization and DNA release upon inflammasome activation (*48-51*). To assess the role of GsdmD in mitochondrial depolarization under the conditions used for this cryo-ET study, we measured the mitochondrial membrane potential of GsdmD^-/-^ iBMDMs upon inflammasome stimulation. Stimulation of GsdmD^-/-^ cells with LPS and nigericin induced depolarization in only 23% of cells (versus 39-48% for WT; **Fig. 5, C** and **D**, and **Fig. S8A** and **B**). Moreover, when LPS and ATP were used for stimulation, WT and GsdmD^-/-^ cells were depolarized to the same extent (**Fig. S8D**). In contrast, knockout of Ninj1, which promotes plasma membrane rupture during pyroptosis (*52*), increased depolarization with either nigericin or ATP stimulation (**Fig. S8, C** and **D**). This suggests that GsdmD contributes to mitochondrial depolarization in cells stimulated with nigericin in a manner independent of plasma membrane rupture. We hypothesized that the GsdmD-dependent component of mitochondrial depolarization could be due to GsdmD pores forming in the outer mitochondrial membrane. Consistent with this, subcellular fractionation experiments showed that upon stimulation with LPS and nigericin GsdmD was proteolytically cleaved and the pore-forming N-terminal proteolytic cleavage fragment was translocated into a purified mitochondrial fraction (**Fig. 5E** and **Fig. S7B**). The outer mitochondrial membrane protein Tom20 was enriched in the purified mitochondrial fraction as expected, but notably, the plasma membrane receptor CD14 was absent in the mitochondrial fraction, indicating that the mitochondrial fraction did not contain any contaminants from the plasma membrane. Cryo-EM images of the purified mitochondrial fraction confirmed that it contained mitochondrial with key morphological features largely preserved (**Fig. S7C**).

## Discussion

The formation of NLRP3 inflammasome signaling assemblies into puncta, or specks, containing ASC and caspase-1 is a cardinal feature of inflammasome activation. Here, our cryo-ET reconstructions, obtained in unstained and fully hydrated conditions, show that the speck is formed of a filamentous network consisting of hollow-tube branched filaments with the dimensions predicted for ASC filaments, based on structural studies of purified ASC PYD filaments and full-length monomeric ASC (*23, 24*). The identification of a filamentous network containing mCerulean-labeled ASC, similar to an ASC punctum seen in a fixed zebrafish section (*26*), positively identifies this protein as the principal filament-forming component in the puncta. These data are supported by analysis of similar structures in wild-type cells labeled with a fluorescent caspase 1 substrate. The filament structure of the ASC with an mCerulean tag on the C-terminus supports the model whereby the PYD forms the filament core, and the CARD decorates the filament. The filament branching visible in the reconstructions is proposed to depend on CARD-CARD interactions because the CARD is required for puncta formation, whereas the ASC PYD alone forms unbranched filaments that lack the structural stability that is inherent to a branched network (*17, 18, 23-26*).

The density of ASC filaments varied in the cryo-ET reconstructions, with multiple core regions of densely packed filaments separated by sparser regions. This suggests that the higher-order structure of the punctum was likely seeded by multiple oligomeric assemblies, rather than growing concentrically from a single nucleation site. In contrast with huntingtin and other neurotoxic protein aggregates, which exclude other macromolecules based on cryo-ET reconstructions (*53*), the structural organization of the ASC filament network allows ribosomes and small Golgi-like vesicles to be retained in the puncta, or to permeate through them.

Permeability of the ASC network to macromolecules could be important for efficient recruitment and release of effector, substrate, and product molecules. Permeability to vesicles would allow active transport of Golgi vesicles carrying activated inflammasome seed-oligomers from sites of NLRP3 activation to the site of punctum formation, as recently proposed (*8*). Moreover, the ribosomes within the puncta could potentially contribute to the inflammasome signaling program, for example by expressing proteins required for signaling, or by responding to danger signals. Indeed, NLRP3 signaling can be activated by translational arrest through direct and indirect mechanisms (*54*), including binding of fungal polysaccharides to ribosomes (*55*).

Our cryo-ET reconstructions, in addition to revealing the novel ultrastructure of ASC within the cell, also allowed morphometric analysis of mitochondria in the inflammasome activated cellular environment. The clearest differences between mitochondria in inflammasome activated cells and unstimulated cells were a two to fourfold reduction in cristae spacing, and the appearance of protein-rich discontinuities of 10-20 nm in diameter in the outer membrane, which are distinct from the larger BAK/BAX macropores (*41, 44*). Permeabilization of both the plasma membrane and outer mitochondrial membrane was reported previously in cells undergoing caspase-1-dependent pyroptosis after inflammasome activation (*42*). Pore formation in the plasma membrane by the N-terminal domain of GsdmD after cleavage by caspase-1 is required for inflammasome-dependent pyroptosis (*46*) and gasdermin D and E pores have been reported to form on the outer mitochondrial membrane following inflammasome activation (*48-51*). The outer-membrane discontinuities in the mitochondria of inflammasome-activated cells had similar dimensions to GsdmD pores, which have an average inner diameter of 22 nm (*46*) (or 13-34 nm based on atomic force microscopy (*47*)). We show here that there is a GsdmD-dependent component to the depolarization mitochondrial membranes in iBMDMs undergoing pyroptosis following stimulation with LPS and nigericin. Moreover, we show that GsdmD is cleaved and the pore-forming N-terminal fragment translocates into the mitochondrial subcellular fraction upon NLPR3 activation. We conclude that GsdmD directly contributes to mitochondrial depolarization during pyroptosis by inserting and forming pores in the outer mitochondrial membrane. Stimulation with ATP instead of nigericin leads to more rapid and extensive loss of mitochondrial potential (**Fig. S8**). The same extensive ATP-induced depolarization occurs in GsdmD^-/-^ and Ninj1^-/-^ cells, suggesting that nigericin and ATP cause depolarization via distinct mechanisms. Further studies are warranted to determine how stimulation with extracellular ATP induces depolarization independent of GsdmD and Ninj1-mediated membrane rupture.

Overall, our structural analyses suggest that the filament branching and packing density within ASC puncta provide structural integrity while allowing downstream signaling molecules to diffuse freely and bind at high density within the network. Our cryo-ET reconstructions provide direct visualization of the cellular organelles (ribosomes and Golgi vesicles) within the ASC speck at a sufficient resolution to support their potential roles in NLRP3 activation. Although many structural details of the NLR inflammasome signaling machinery remain unclear, this study demonstrates the potential for combined cryo-CLEM and cryo-ET approaches to extract detailed, hypothesis-generating ultrastructural information for fully assembled innate immune signaling complexes in their cellular context.

## Supporting information

Combined Supplementary Information

## Acknowledgments

We thank Dustin Morado for assisting in cryo-ET data collection and image processing. We thank Camilla Ventura Santos and Kunimichi Suzuki (MRC-LMB) for advice on subtomogram averaging with WARP. We thank Wanda Kukulski, Emma Jones, Robert Pickering, Maria Daly, Victoria L. Hale, Zunlong Ke, and Panagiotis Tourlomousis for advice and helpful discussions. We acknowledge the following core facilities at the MRC Laboratory of Molecular Biology for access, training, and support: Electron Microscopy, Flow Cytometry, Light Microscopy, and Scientific Computing.

## Author contributions

Conceptualization: YL, LJH, CEB, YM

Formal Analysis: YL, HZ, HA, JB, JDH, KEL

Methodology: YL, CEB, YM

Investigation: YL, HZ, JH, ACB, CH, LJH

Visualization: YL, HA, YM

Funding acquisition: CEB, YM

Project administration: CEB, YM

Supervision: CEB, YM

Writing – original draft: YL, CEB, YM

Writing – review & editing: YL, CEB, YM

## Data and materials availability

Representative electron tomograms were deposited in the Electron Microscopy Data Bank (EMDB) with accession codes EMD-13585 and EMD-13586. The scripts used in this study and instructions for use are available from Github: https://github.com/DustinMorado/subTOM, https://github.com/jboulanger/imagej-macro/tree/main/FRAP_Measure, https://github.com/alisterburt/dynamo2m, https://github.com/builab/subtomo2Chimera.

## Supplementary Materials

Materials and Methods

Figs. S1 to S8

References (*56-65*)

Movies S1 to S4

Data S1 to S4

