## Supplementary material for "Cryo-electron tomography of NLRP3-activated ASC complexes reveals organelle co-localization": Combined Supplementary Information

**This PDF file includes:**

Materials and Methods
Figs. S1 to S8
References (*56–65*)
Captions for Movies S1 to S4
Captions for Data S1 to S4

**Other Supplementary Materials for this manuscript include the following:**

Movies S1 to S4
Data S1 to S4

### Materials and Methods

Mammalian cell culture

Wild-type immortalized primary mouse bone-marrow derived macrophages (WT iBMDMs) and ASC-mCerulean iBMDMs, kindly provided by Eicke Latz (Univ. Bonn), were cultured in Dulbecco's modified Eagle Medium high glucose (DMEM; Gibco), supplemented with 10% v/v foetal bovine serum (FBS; Gibco). Human monocytic THP-1 cells (European Collection of Authenticated Cell Cultures) were grown in Roswell Park Memorial Institute (RPMI) 1640 Medium (Gibco), supplemented with heat inactivated 10% v/v FBS, 10 mM HEPES pH 7.4, 1 mM Sodium Pyruvate and 0.05 beta-mercaptoethanol. Cells were checked regularly for mycoplasma contamination by MycoAlert™ mycoplasma detection kit (Lonza) and cells were free of mycoplasma contamination.

Inflammasome stimulation

For stimulation of WT iBMDMs and ASC-mCerulean iBMDMs, cells were primed with 200 ng/ml LPS for 3 h in Opti-MEM® (Gibco) or DMEM supplemented with 10% FBS before NLRP3 stimulus was used. The following stimulus were used: 10 μM nigericin (Sigma or Invivogen) and 5 mM ATP (pH adjusted to 7.4; Sigma) for 30 min or 60 min with pan-caspase inhibitor Z-VAD-FMK (Invivogen) or carboxyfluorescein-labelled inhibitor of caspase-1 YVAD (FAM-FLICA; ImmunoChemistry Technologies) for *in situ* cryo-ET sample preparation unless otherwise specified. For stimulation of THP-1 cells, priming and stimulation were performed in RPMI Medium and conditions were listed above.

Cytokine measurement and immunoblotting

IL-1β secretion was measured by ELISA as previously described (*27*). Briefly, supernatants were collected after inflammasome activation and the OptEIA kit (BD BioSciences) was used to measure mouse IL-1β concentration in cell supernatant according to the manufacturer's instructions. For the immunoblot in **Fig. S1**, ASC-mCerulean iBMDMs were lysed in cell lysis buffer (150 mM NaCl, 50 mM Tris-HCl pH 8, 1% Triton X-100, 1 mM PMSF, 10 μg/mL leupeptin, 1 μg/mL aprotinin) on ice for 10 min before centrifugation. Supernatant was collected and boiled in SDS sample loading buffer for 5 min. Following separation by 4-20% gradient SDS-PAGE, the proteins were transferred to a PVDF membrane. Membrane was blocked in 5% milk in PBS + 0.2% Tween-20. The primary antibody was rabbit monoclonal anti-mouse GsdmD (Abcam, ab209845, RRID:AB_2783550). The secondary antibody was goat anti rabbit IgG-HRP (Santa Cruz Biotechnology, sc-2004, RRID:AB_631746).

Immunofluorescence staining

For immunofluorescence, cells were plated as monolayer on μ-Slide 8-Well chamber (ibiTreat, Ibidi, 80826) or 12-well chamber (removable, Ibidi, 81201). Cells were primed and stimulated as above, and then fixed with 4% paraformaldehyde for 5 to 10 min at room temperature. Cells were permeabilised and blocked in 0.1% Saponin (Sigma, 47036) supplemented with 20% FBS (Gibco) or 2% Bovine Serum Albumin (Sigma, A7030) in PBS with primary antibody overnight. Washing was performed before cells were incubated with secondary antibodies. The following primary antibodies were used: rabbit anti-ASC rabbit polyclonal pAb AL177, 1:200 dilution (AdipoGen, AG-25B-0006, RRID:AB_2490440); rabbit anti-ASC monoclonal ASC/TMS1(D2W8U), 1:800 dilution (Cell Signalling Technology, 67824,

RRID:AB_2799736); goat anti-NLRP3 polyclonal, 1:200 dilution (Abcam, ab4207, RRID:AB_955792); goat anti-IL-1β polyclonal, 1:500 dilution (R&D Systems, AF-401-NA, RRID:AB_416684); mouse anti-γ-tubulin monoclonal, 1:400 dilution (Sigma-Aldrich, T6557, RRID:AB_477584); rabbit anti-TGN38 polyclonal, 1:250 dilution (Novus Biologicals, NBP1-03495, RRID:AB_1522533); rabbit anti-TOM20 FL-145 polyclonal, 1:80 dilution (Santa Cruz Biotechnology, sc-11415, RRID:AB_2207533). The Alexa Fluor secondary antibodies (488, 555, 568 and 647) used were: 488-labelled donkey anti-goat IgG (H+L) (1:500, ThermoFisher, A11055, RRID:AB_2534102); 555-labelled goat anti-rabbit IgG(H+L) (1:500, ThermoFisher, A21428 RRID:AB_141784); 568-labelled goat anti-mouse IgG (H+L) (1:500, ThermoFisher, A11004, RRID:AB_2534072); 647-labelled goat anti-rabbit IgG (H+L) (1:500, ThermoFisher, A21244, RRID:AB_2535812). After incubation with secondary antibodies at room temperature for 1 h, cells were washed three times with blocking buffer, one time with PBS and final wash with water before mounting with ProLong Gold Antifade Mountant with DAPI (ThermoFisher).

Confocal fluorescence imaging

Single point-scanning confocal microscopy was carried out on a Zeiss (Oberkochen, Germany) LSM 780 or LSM 710 microscope using a 40x/1.3 NA Fluar or 63x/1.4 NA Plan-apochromat oil immersion objective lens. The microscope was equipped with 405, 458, 488, 514, 561 and 633 nm laser lines. Multi point-scanning confocal microscopy was carried out on Visitech (Sunderland, UK) iSIM mounted on a Nikon (Tokyo, Japan) Ti2 microscope stand using a 100x/1.49 NA SR Apo TIRF oil immersion objective lens or a 60x/1.2 NA Plan Apo VC water immersion objective lens. The iSIM was equipped with 405, 445, 488, 561 and 640 nm laser lines and Hamamatsu (Hamamatsu, Japan) ORCA-Flash4.0 V3 sCMOS cameras. Filter ranges: green (500-545 nm), red (593-624 nm). Live-cell samples were heated to 37˚C and supplemented with 5% $CO_2$ using a microscope incubation chamber.

Live-cell staining and Fluorescence Recovery after Photobleaching (FRAP)

For live-cell staining of activated caspase-1, the supernatant was removed following inflammasome stimulation and replaced with DMEM supplemented with 10% (v/v) heat-inactivated FBS containing 0.5x reconstituted FAM-FLICA (ImmunoChemistry Technologies). For live-cell Golgi staining, LPS-primed iBMDMs were labelled with BODIPY TR ceramide (ThermoFisher, D7540) according to the manufacturer's protocol, prior to stimulation with nigericin. Cells were imaged as described above.

To quantify fluorescence of FAM-FLICA, BODIPY TR ceramide and ASC-mCerulean in the speck-forming area of a cell following stimulation (**Fig. 3A**), the cell outline was traced at each timepoint with the script Morph_ROI.ijm (https://github.com/jboulanger/imagej-macro/blob/main/FRAP_Measure/Morph_ROI.ijm). Fluorescence intensity within the speck-forming area and whole-cell fluorescence intensity and were measured at different time points. The graphs in **Fig. 3A** report what percentage of the whole-cell fluorescence mapped to the area of speck formation at two timepoints: at the beginning of the movie (1-3 min after addition of nigericin) and at the time when speck formation reached completion (20-50 min after addition of nigericin).

For confocal FRAP, ASC-mCerulean iBMDMs were primed and induced as described above with the presence of caspase-1 inhibitor (Z-VAD-FMK). Live-cell FRAP imaging was performed on a Zeiss LSM 710 microscope equipped with a 63x/1.4 NA Plan-apochromat oil immersion objective lens and a 458 nm laser line for excitation and bleaching. The sample environment was heated to 37˚C and supplemented with 5% $CO_2$ using a microscope incubation chamber. ASC-mCerulean speck was photobleached with 100% laser power. Images were acquired at 3 s intervals. Images were collected at three pre-bleach timepoints and for 160 s post-bleaching. Movies were analysed in ImageJ/Fiji (*43*) using customised script FRAP_measure.ijm (https://github.com/jboulanger/imagej-macro/blob/main/FRAP_Measure/FRAP_measure.ijm). Briefly, measures were normalised to account for the general photobleaching caused by image acquisition as well as sample motion over time. Normalised fluorescence intensity measurements were obtained after background and bleaching corrections. The timeframe varied for experiments (n = 8). Each measurement was interpolated using the normalised fluorescence intensity measurements and imported to Python for plotting of the fluorescence intensity curve.

Quantification of mitochondrial membrane potential

After 30 min of inflammasome stimulation as described above, iBMDMs were incubated with 20 nM TMRM (Life Technologies) for 15 min. iBMDMs were incubated for 3 h in 50 µM of the mitochondrial uncoupler carbonyl cyanide 3-chlorophenylhydrazone (CCCP; ThermoFisher, M20036). Control samples included: untreated; TMRM+; TMRM+CCCP+; TMRM+LPS+ iBMDMs. TMRM fluorescence was quantified by flow cytometry and data were acquired on Eclipse flow cytometer (Sony Biotechnology). Quantification was set at 100,000 for cell count and a spectrum window of FL3 (595BP) was used to detect TMRM signal. Data were analysed and visualised using FlowJo10.

Cryo-ET sample preparation

Quantifoil Au 200-mesh finder grids (R2/1 or R2/2, Quantifoil Micro Tools) were glow discharged with a 30 mA current for 30 s with an Edwards S150B Sputter Coater. Grids were sterilized by UV irradiation for 10 min and immersed in PBS supplemented with 10 µg/ml fibronectin (Sigma) overnight in 8-well or coculture wells (ibidi). Grids were washed with PBS three times. Cells were seeded in 8-well or coculture wells (ibidi) and incubated overnight at 37˚C and 5% $CO_2$. Cells cultivated on grids were primed, induced with inflammasome stimuli and were plunge-frozen in liquid ethane using Leica EM GP2 cryo-plunger. Prior to plunging, 4 µl of cell culture medium was added to the cell side and backside blotting was applied for 6-8 s. The chamber conditions were maintained at 37˚C, 100% humidity during freezing. Grids were stored in liquid nitrogen.

Cryo-fluorescent light microscopy (Cryo-FM) and cryo-focused ion-beam (FIB) milling

Grids were screened for cells with ASC/caspase-1 speck by light microscopy using a Leica EM Cryo CLEM microscope equipped with a cryo-stage, an ORCA-Flash4.0 V2 sCMOS camera (Hamamatsu Photonics) and a 50x/0.9 NA HCX PL APO cryo-objective lens. Montage acquisition of grids was performed with Leica LAS X software, while recording the following channels: green (L5 filter, 50 ms), far red (Y5 filter, 20 ms), and brightfield (50 ms). Z-stacks were recorded at 0.5 µm with step size 21 for each grid square to determine the best focus for the montage. The montage was completed by stitching the best focus image from the Z-stack with Leica LAS X.

Lamellae were prepared using a Scios Dual Beam FIB scanning electron microscope (SEM; ThermoFisher) equipped with a Quorum PP3010T cryo-stage. The milling protocol was adapted from a previously published method (*29*). Grids were coated with organometallic platinum using the gas injection system (GIS; ThermoFisher) operated at RT, for 8 s, at 12 mm working distance and 25˚ stage tilt. A first rough milling was performed at 25˚ stage tilt with a 30 kV ion beam voltage and 1 nA current until the lamella thickness reached 10 µm. The micro-expansion joints were applied to improve lamella stability (*32*) using the milling parameters listed above. The stage was tilted to 20˚ for subsequent milling steps. Rough milling steps were applied as following with 30 kV ion beam voltage: a 5 µm lamella thickness was reached with current 0.5 nA; 3 µm lamella thickness was reached with a 0.3 nA current; and 1 µm lamella thickness was obtained with a 0.1 nA current. Fine milling to a final lamella thickness of approximately 200 nm was performed with ion beam settings of 30 kV and 50 pA or 10 pA, or 10 kV and 23 pA or 11 pA.

Grids were loaded on a Linkam CMS196V cryo-stage, and lamellae were imaged on a Zeiss microscope equipped with Axiocam 503 mono and Colibri 7-illumination module R(G/Y) CBV-UV. Z-stacks of the lamellae were acquired to identify ASC/caspase-1 specks using a 200-300 nm step size. Only 3% to 6% of the lamellae (one lamella out of 17-29 lamellae) retained fluorescent signal from ASC-mCerulean or FAM-FLICA after milling. A projected cryo-fluorescent image was used to correlate with an SEM image using the eC-CLEM plugin from ICY (*33*). Registration between the lamella map and SEM image was achieved by transforming coordinates using edges or salient features of the lamella. The maximum projection Z-stack was then saved for correlation with SerialEM (*32*,*33*).

Tilt-series acquisition and tomogram reconstruction

Grids with a fluorescent lamella were transferred to a 300 kV Titan Krios electron microscope (ThermoFisher) equipped with an energy filter (Gatan). Movies were acquired with a K2 or K3 direct electron detector with SerialEM (*34, 35*). Cryo-FM/EM correlation was then used to locate an area of interest for tilt-series acquisition. Briefly, a medium-magnification montage EM map (MMM) of the lamella was generated and a fluorescence map (FM) was loaded into SerialEM. The maps were correlated through recognition of geometric edges of the lamella. Transformation of coordinates yielded an overlay map which provided guidance for tilt-series acquisition. Tilt-series were collected at a nominal 42,000X, 33,000x or 26,000x magnification, resulting in pixel size 2.13 Å, 2.69 Å or 3.42 Å, over a tilt range of -60˚ to +60˚ with 1˚, or 2˚ increments, a total dose of 140-240 electrons $A^{-2}$ and a nominal defocus range of -4 to -8 µm. A dose-symmetric scheme was used (*56*). A pre-processing script was used from SubTOM (by Dustin Morado; https://github.com/DustinMorado/subTOM). Frames were aligned using IMOD (*36*). Reconstruction was performed by weighted back-projection, and segmentation with IMOD. The MATLAB script deconv from Warp (*57*) was used for visualisation. Filament tracing was performed manually with IMOD and UCSF Chimera (*58*). Membrane segmentation was performed with TomoSegMemTV (*59*).

Subtomogram averaging of ribosomes

302 ribosomes were manually picked in Dynamo (*60*). Coordinates were converted to Warp format with the script dynamo2warp.py (https://github.com/alisterburt/dynamo2m/blob/master/dynamo2m/dynamo2warp.py).

Subtomograms and corresponding contrast transfer function (CTF) models were reconstructed in Warp (*57*) with a box size of 44 pixels, a pixel size of 12 Å, and a particle diameter of 350 Å for normalization. Initial subtomogram alignment and averaging was performed in RELION v3.1 (*61*) using a previously determined *in situ* cryo-ET structure of the mammalian 80 *S* ribosome (*62*), low pass-filtered to 60 Å resolution, as the reference. This subtomogram average was used to perform template matching in Warp at 10 Å per pixel, with the addition of 808 particles. False positives were removed from template matching results. Subtomograms were exported from Warp to RELION for 3-D classification. A 3-D class with 1,058 particles was selected without alignment and reference. 3-D refinement was performed on the selected particles. The refined positions file was converted to RELION3.0 format with the script relion_star_downgrade.py (https://github.com/alisterburt/dynamo2m/blob/master/dynamo2m/relion_star_downgrade.py). Particles were exported at with pixel size of 6 Å and refined using the average density as the reference. The resolution of the subtomogram average was calculated to be 23 Å during postprocessing in RELION v3.1 (using a Fourier shell correlation cut-off of 0.143). Ribosome subtomogram average volumes were mapped back into the 3-D segmented model with the script relionsubtomo2ChimeraX.py (https://github.com/builab/subtomo2Chimera/blob/main/relionsubtomo2ChimeraX.py).

Subcellular fractionation and immunoblotting

Mitochondrial and cytosolic fractions were isolated using the Mitochondria/Cytosol Fractionation Kit (Abcam, ab65320). Briefly, 300 mg of ASC-mCerulean iBMDMs and WT iBMDMs were harvested after stimulation with LPS and nigericin. All centrifugation steps were performed at 4˚C. Cells were washed with ice-cold PBS and pelleted at 600 g for 5 min and resuspended in 1 ml 1x Cytosolic Extraction Buffer without protease inhibitors. After 10 min on ice, cells were homogenized on ice with 100 passes on pestle B. The cell lysate was centrifuged at 700 g for 10 min. The pellet ("PM pellet" in **Fig. 5** and **Fig. S7**), containing plasma membrane and any unlysed cells, was resuspended in RIPA buffer (50 mM Tris, 150 mM NaCl, 1% Triton X-100, 1 mM EDTA, 0.1 % SDS). The supernatant was collected and centrifuged at 10,000 g for 30 min. The resulting supernatant was the cytosolic fraction ("Cytosol" in **Fig. 5** and **Fig. S7**). The pellet was resuspended in ice-cold 20 mM HEPES-KOH pH 7.5, 250 mM sucrose, 1 mM EDTA, and loaded onto a sucrose gradient prepared as described (*63*). Briefly, the sucrose gradient was prepared by placing 1.5 ml of 60% sucrose buffer (60% sucrose, 20 mM HEPES-KOH pH 7.4, 1 mM EDTA) into an SW40 centrifuge tube, followed by 4.5 ml 32% sucrose buffer, 1.5 ml 23% sucrose buffer, and 1.5 ml 15% sucrose buffer. The suspension was then centrifuged in an SW40Ti rotor at 28,000 rpm for 1 h at 4˚C. A brown band containing the mitochondrial fraction ("Mitoch." In **Fig. 5** and **Fig. S7**) formed at the boundary between the 32% and 60% sucrose solutions and was extracted with a pipette. The cytosolic, mitochondrial and pellet fractions were immediately incubated at 95˚C for 10 min in 2x SDS-PAGE loading buffer and used for immunoblotting or stored at -20˚C.

The primary antibodies used for immunoblotting were as follows. Rabbit monoclonal anti-mouse GsdmD [EPR19828], 1:1,000 dilution (Abcam, ab209845, RRID:AB_2783550); plasma membrane marker: rabbit monoclonal anti-mouse CD14, 1:1,000 dilution (Abcam, ab221678, RRID:AB_2935854); cytosolic marker: mouse monoclonal anti-GAPDH, 1:5,000 dilution (Proteintech, 60004-1-Ig, RRID: AB_2107436); mitochondrial marker: rabbit polyclonal anti-TOM20 FL-145, 1:500 dilution (Santa Cruz Biotechnology, sc-11415, RRID:AB_2207533).

Quantifications and statistical analysis

Filament lengths were measured using IMOD (*36*) from five tomograms of ASC-mCerulean iBMDMs and one tomogram of WT iBMDMs labelled with FAM-FLICA. Histograms were plotted with MATLAB. Branching angles were measured with Fiji (*43*) and exported to MATLAB for plotting. Similarly, diameters of trans-Golgi-like vesicles were measured in Fiji and exported to MATLAB for plotting.

The ASC filament core diameter was determined from 8 Å-thick virtual tomographic slices from the ASC-mCerulean tomogram following neural network image restoration with cryoCARE (*38*). Density line profiles 30 pixels in length (8 Å/pixel) were recorded perpendicular to the ASC filaments in Fiji. The density maxima corresponding to the core tube walls were identified in the density profiles (n =13 for ASC-mCerulean) by calculating the first derivative of the smoothed density profile curve in GraphPad Prism v9.5.1 (the maxima were identified in the first derivative curve as x-axis intercepts with a negative slope). The density profiles were aligned by centring them on the midpoint of their respective tube-wall maxima positions. The aligned profiles were superimposed and used to fit a "sum of two Gaussians" function by non-linear regression in GraphPad Prism. The separation between the two maxima in the resulting function was taken as the filament core diameter.

Mitochondrial tomograms were loaded in Fiji (*43*) and ROI manager was used to measure morphological parameters, which were compared using Two-way ANOVA and plotted with GraphPad Prism. For the inner-to-outer membrane spacing, the distance between the nearest edges of the inner and outer membranes was measured. The cristae width was measured between the inner membranes of the cristae. The spacing between cristae were measured using the perpendicular distance between the internal border of the neighbouring cristae. The inner-to-outer membrane spacing, cristae width and inter-cristae spacing were measured perpendicular to the membranes, evenly spaced for each tomogram, with thirty values per slice, for four slices per tomogram, from eight tomograms. The cristae apex angles were measured for every tip visible in ten slices per tomogram. Every value was plotted along with the average. If the end of a crista was flat, then the measurement of the two angles on either side was measured. Any measurement was only taken if the internal border of the membrane was clearly distinguishable from the matrix. All measurements were evenly spaced through each slice, to give a value representative of the whole slice, using all the mitochondria present in each image. All measurements were taken using 8 different tomograms: four control, two 30-min nigericin-treated, and two 60-min nigericin-treated mitochondria.

For measurement of the outer-mitochondrial membrane gap density, we calculated the density ratio as follows. A total of 21 virtual slices were selected in a region containing a visible outer-membrane gap. The average pixel grey value was calculated from circular areas 13.6 nm in diameter: three cytosolic areas, three inter-membrane space areas and an outer-membrane gap area. This was repeated for each of the 21 virtual slices. The ratio of average outer-membrane gap site to cytosolic grey value, and the ratio of average inter-membrane space site to cytosolic grey value were calculated for each virtual slice. The grey area ratio was then plotted with Igor (WaveMetrics, Inc.) to show the change of protein density in Z direction.

Correlative fluorescence microscopy and electron tomography of resin-embedded cells

Correlative fluorescence microscopy and electron tomography (RT-CLEM) of resin-embedded cells was performed as described (*41,64*). Briefly, ASC-mCerulean cells were grown on carbon-coated 3 mm sapphire discs (Wohlwend GmbH) in two-well chambers (ibidi) for 24 h before priming and NLRPC-driven inflammasome stimulation as described above. Cells were stained with MitoView far-red and FLICA before high pressure freezing with an HPM100 high pressure freezing system (Leica Microsystems). Grids were imaged with a Leica EM Cryo CLEM cryo-fluorescence microscope to localize cells containing ASC-mCerulean specks. Freeze substitution was preformed using 0.008% uranyl acetate in acetone and embedded in Lowicryl HM20 (Polysciences) using an AFS2 (Leica Microsystems). Blocks were sectioned into 300 nm thin sections using a microtome (Leica Microsystems) equipped with a diamond knife (Diatome). The sections were collected on 200 mesh/300 mesh copper grids with carbon support (Agar Scientific). TetraSpeck 100-nm microspheres were diluted 1:100 in PBS and applied to the sections for use as fiducial markers for correlation. Sections were imaged on a Nikon Ti2 wide field microscope equipped with a Niji LED light source (Bluebox Optics), a x100/1.49 NA Apo TIRF oil immersion objective lens and a Neo sCMOS DC-152Q-C00-FI camera (Andor Technology). Filters: mCerulean (89006 filter set, Chroma Technology), fluorescein (49002 filter set; Chroma Technology), MitoView Far Red (49006 filter set; Chroma Technology). EM images were collected using a Tecnai F20 electron microscope (ThermoFisher) operated at 200 kV and a high-tilt tomography holder (Fischione Instruments, Model 2020). An image montage of regions of interest was acquired on a BM-Orius detector using TEM mode at 150-200 µm defocus using SerialEM v3.8.0 (*35*) at a pixel size of 1.1 nm. The correlation between the montaged map and fluorescent map was performed by image transformation of registered fiducial markers in both image modalities as previously described (*65*). Tilt-series were acquired from approximately -60 to +60 with 1˚ increment at a pixel size of 1.1 nm. Samples were rotated 90˚ to acquire dual axis tilt-series. Tomograms were reconstructed and visualized with IMOD (*36*).

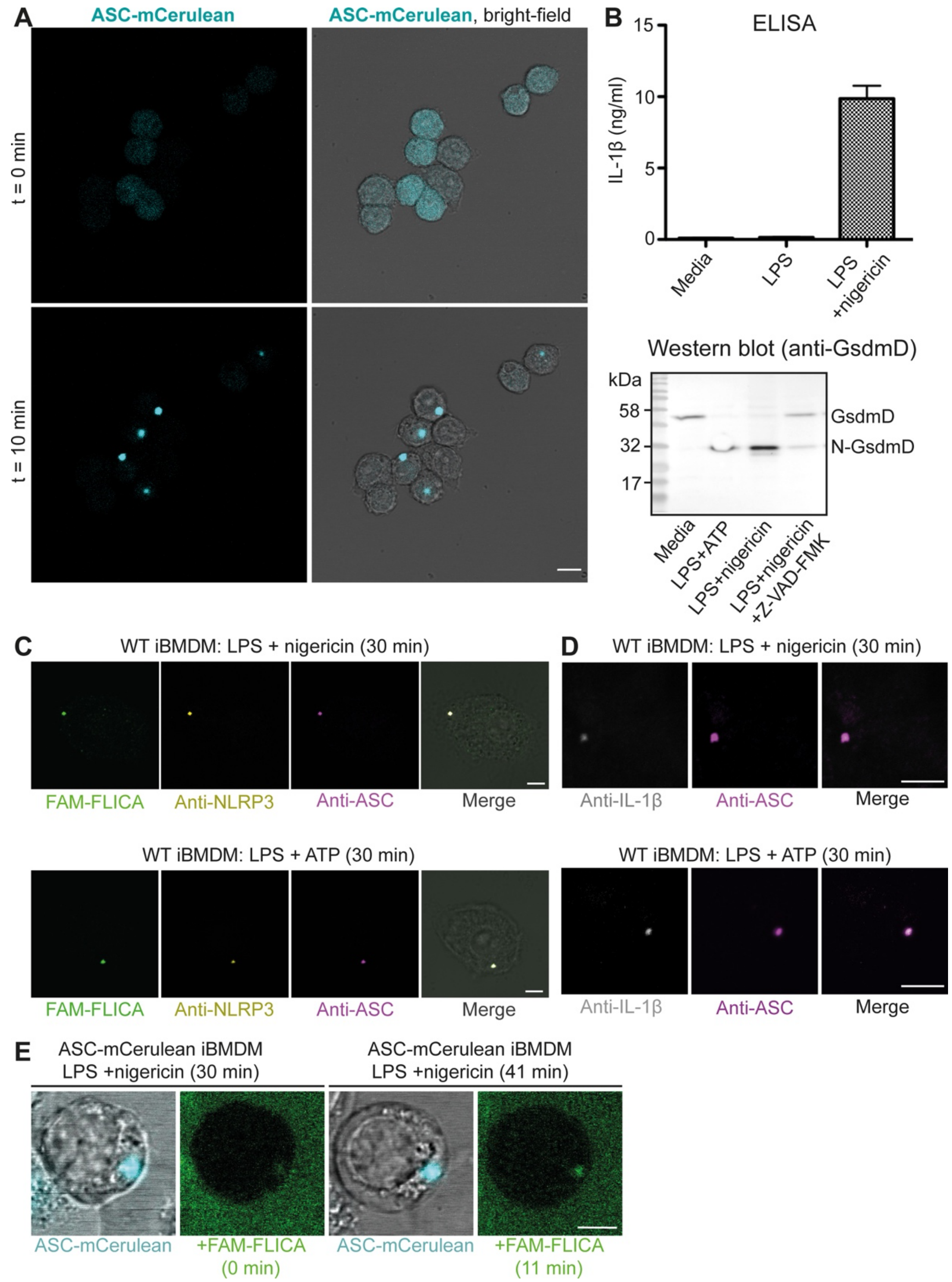


**Fig. S1. Tracking NLRP3 inflammasome activation in iBMDMs with fluorescence confocal microscopy and biochemical assays.** (**A**) Fluorescence and bright-field micrographs of live iBMDMs expressing ASC-mCerulean. ASC specks formed within 10 min of NLRP3 activation with LPS and nigericin. Scale bar, 10 µm. (**B**) Upper panel, ELISA measuring IL-1β secretion in NLRP3-activated ASC-mCerulean iBMDMs; lower panel, Western blot showing proteolytic cleavage of GsdmD to yield the pore-forming N-terminal fragment (N-GsdmD). (**C**) Fluorescence micrographs of WT iBMDMs stimulated with nigericin or ATP. Caspase-1 was detected with FAM-FLICA. NLRP3 and ASC were detected by immunostaining. Scale bars, 5 µm. (**D**) Immunofluorescence micrographs of WT iBMDMs stimulated with nigericin or ATP, stained for IL-1β and ASC. Scale bars, 5 µm. (**E**) Snapshots of live nigericin-stimulated ASC-mCerulean iBMDMs following addition of FAM-FLICA. Scale bar, 5 µm.

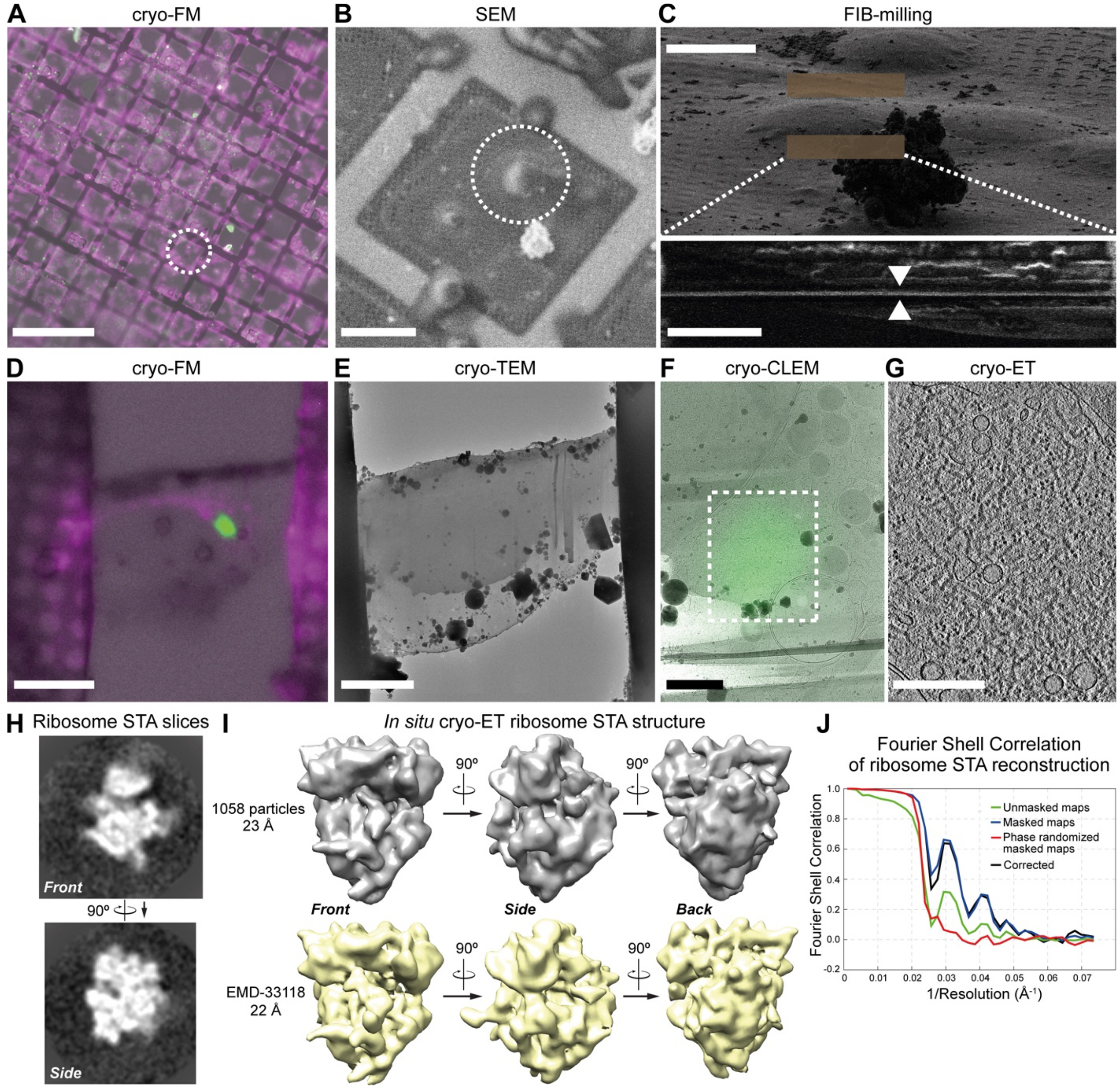


**Fig. S2. Workflow for correlative cryo-light and electron microscopy (cryo-CLEM).** (**A**) Cryo-fluorescence microscopy (cryo-FM) of ASC-mCerulean iBMDMs, with brightfield image overlaid. Cells were cultured on finder grids, primed with LPS, stimulated with nigericin for 30 min, labeled with MitoView and FAM-FLICA, vitrified, and imaged. The white circle denotes an area of interest. Scale bar: 300 µm**.** (**B**) Grids were transferred to a scanning electron microscope (SEM) fitted with a focused-ion-beam (FIB). The cell containing the area of interest, identified using the finder-grid markers, was imaged by SEM. Scale bar, 40 µm**.** (**C**) FIB-induced secondary electron images of the area shown in (B) before and after FIB milling: Upper panel, before FIB milling with initial target milling windows shown in brown (scale bar, 10 µm); lower panel, side-on view of the lamella after FIB milling (scale bar, 4 µm). (**D**) Cryo-FM image of a lamella containing an ASC speck with brightfield image overlaid. Scale bar, 10 µm. (**E**) Transmission cryo-EM map of the lamella shown in (D). Scale bar, 7 µm. (**F**) Cryo-EM map overlaid with the cryo-FM map after registration and transformation. The region of interest for tilt-series acquisition is boxed. Scale bar, 1 µm. (**G**) Reconstructed tomographic slice acquired in the area boxed in (F). Filaments and vesicles are visible. Scale bar, 300 nm. See **Movie S2** for a full tomogram. (**H**) Slices through a 23 Å resolution cryo-ET subtomogram averaging (STA) reconstruction of 1058 ribosomes from six tomograms of ASC-mCerulean iBMDMs. (**I**) Upper panels, volume representations of the STA reconstruction shown in (H). Lower panels, *in situ* STA volumes of ribosomes from Ref. (*62*) with a 22 Å low-pass filter applied. (**J**) Fourier shell correlation plots for the ribosome STA reconstruction.

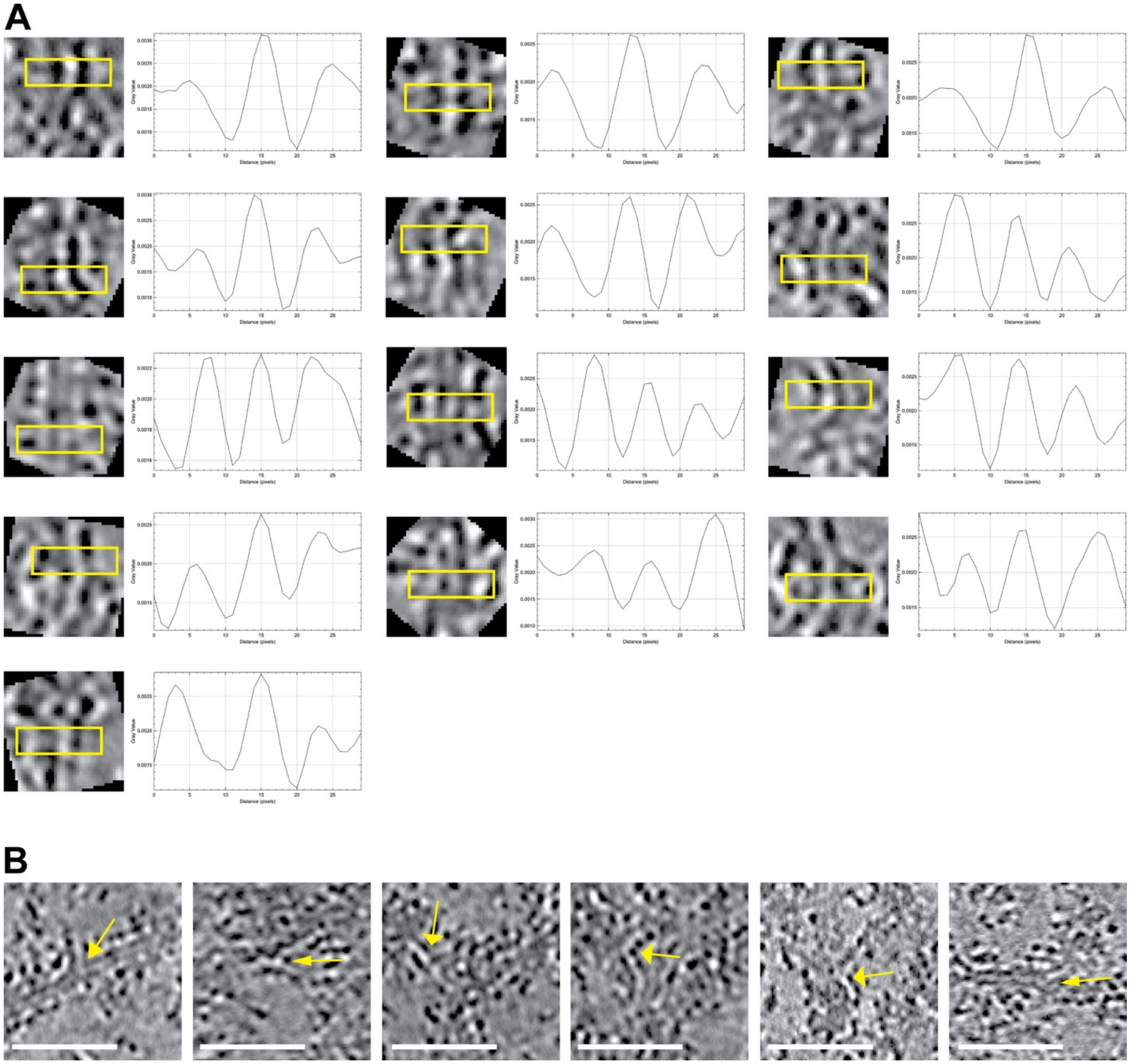

**Fig. S3. Measurement sites for ASC filament properties on cryo-ET tomograms.** (**A**) The thirteen 8-Å thick tomogram slices and associated density profiles plots that were used to measure the ASC filament core diameter (see **Fig. 2, B** and **C**). The tomogram slices were restored with cryoCARE (*38*). 30-by-8-pixel (24 x 6.4 nm) cross-section areas, boxed in yellow, were drawn in Fiji (*43*) perpendicular to the ASC filaments. Density profiles were plotted from these areas in Fiji with the Plot Profile function. The pixel size is 8 Å. (**B**) Tomogram slices showing filament branching angle measurement sites, marked by yellow arrows. Scale bars, 50 nm.

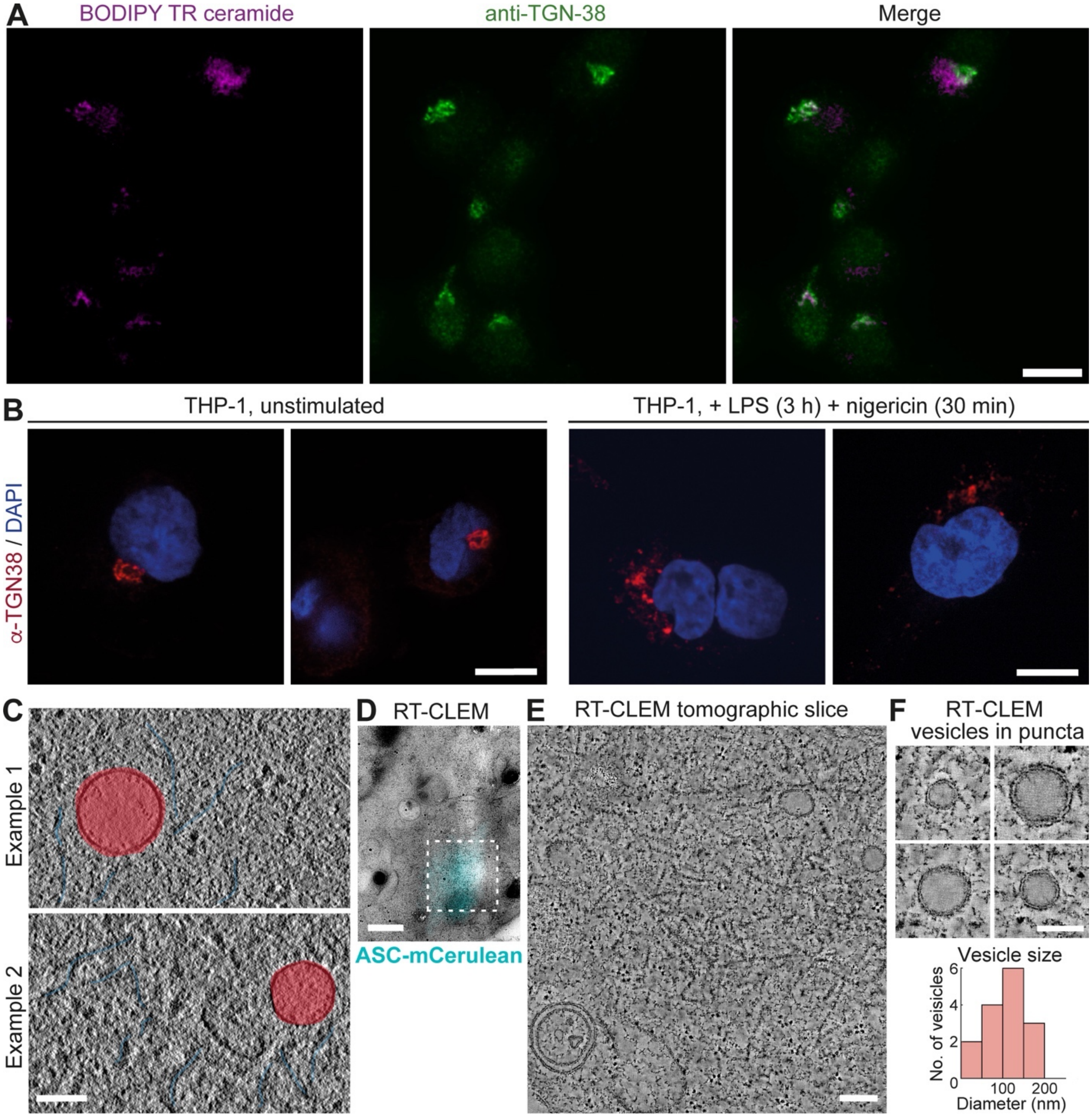


**Fig. S4. Imaging of Golgi-like vesicles within ASC puncta.** (**A**) Unstimulated WT iBMDMs stained with BODIPY TR and anti-TGN38 antibody. Scale bar, 10 µm. (**B**) THP-1 cells stained with anti-TGN38 antibody. Scale bars, 10 µm. (**C**) Tomographic slices from an ASC-mCerulean speck showing vesicles adjacent to ASC filaments. Scale bar, 100 nm. (**D-F**) ASC-mCerulean iBMDMs stimulated with nigericin for 30 min, high-pressure frozen, chemically fixed, sectioned and imaged by CLEM at room temperature. (**D**) Low magnification EM map overlaid with ASC-mCerulean fluorescence. Scale bar, 1 µm. (**E**) 5.5 nm-thick tomographic slice of the region boxed in (D). Scale bar, 200 nm. (**F**) Tomographic slices showing vesicles with low-density lumens within ASC-mCerulean puncta. Scale bar, 100 nm. The histogram shows the vesicle diameter distribution.

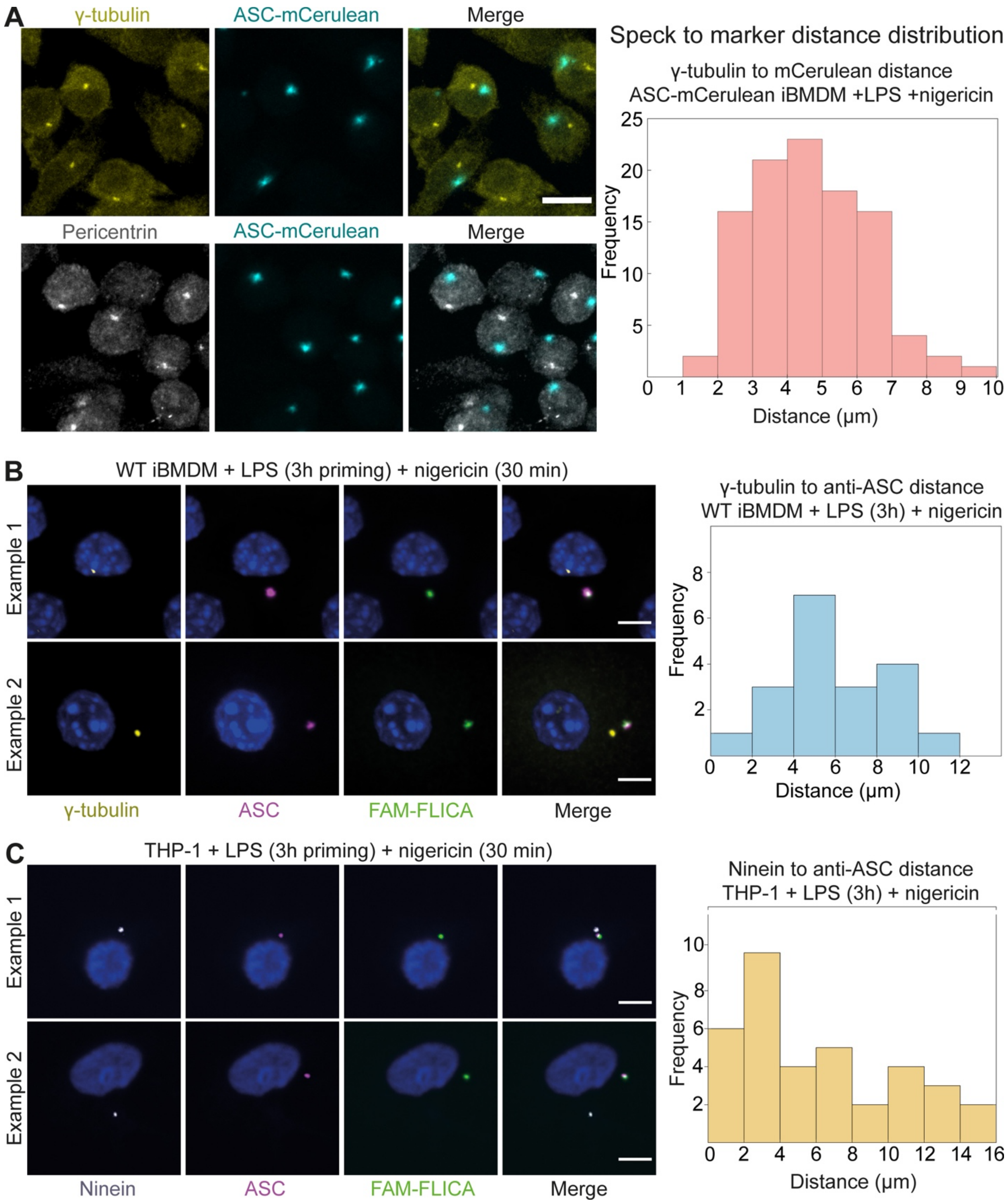


**Fig. S5. Localization of ASC and caspase-1 relative to MTOC-associated components.** (**A**) Fluorescence confocal micrographs of ASC-mCerulean iBMDMs stained with antibodies against γ-tubulin or pericentrin. Right, distance distribution between ASC-mCerulean and anti-γ-tubulin fluorescence foci. (**B**) Fluorescence micrographs of WT iBMDMs stained with FAM-FLICA and antibodies against γ-tubulin and ASC. Right, distance distribution between ASC/FAM-FLICA and anti-γ-tubulin fluorescence foci. (**C**) Fluorescence micrographs of THP-1 cells stained with FAM-FLICA and antibodies against ninein and ASC. Right, distance distribution between ASC/FAM-FLICA and anti-ninein fluorescence foci. Scale bars, 10 µm.

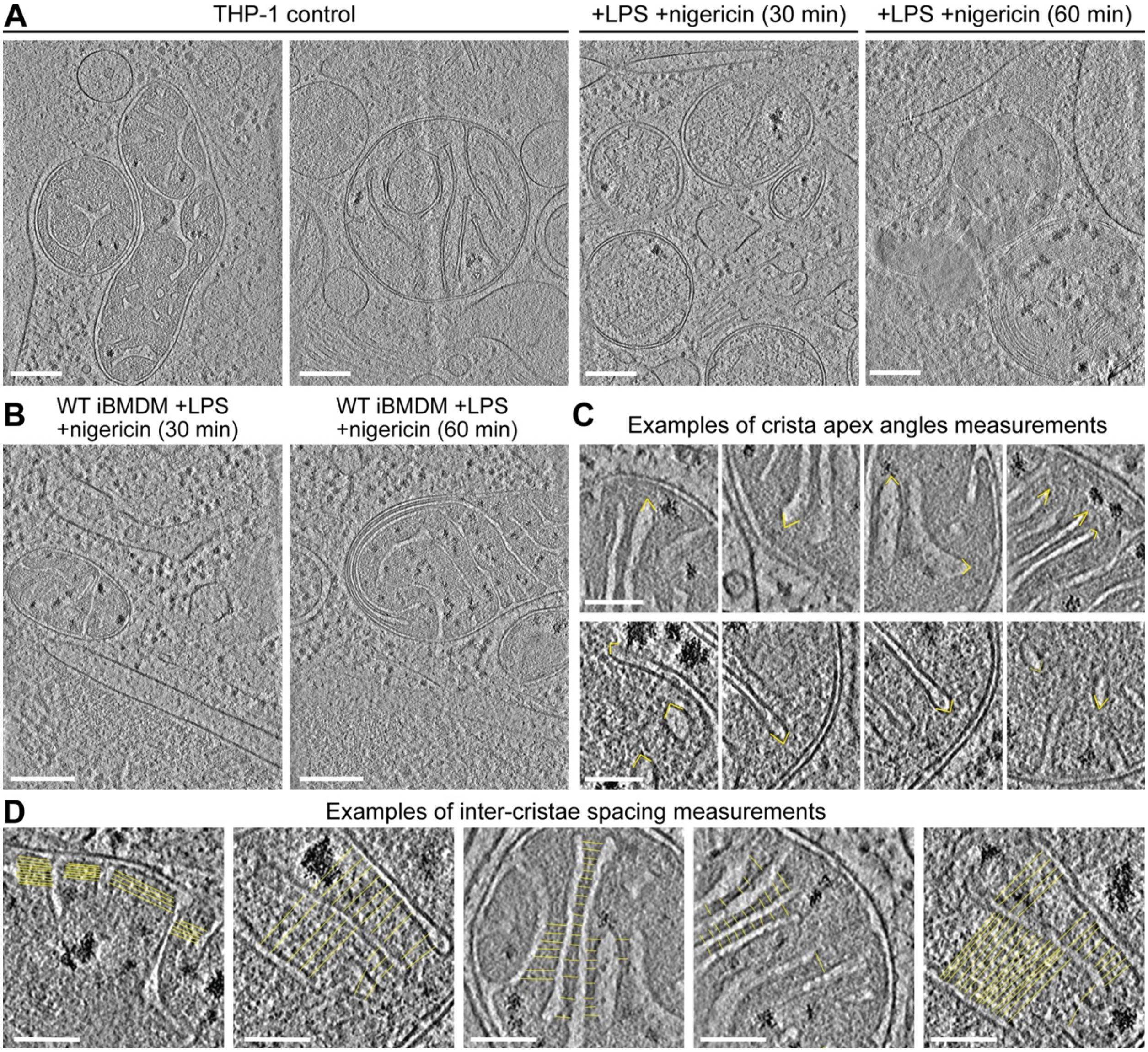


**Fig. S6. Mitochondrial morphology of NLRP3-activated THP-1 cells and quantification of mitochondrial morphology.** (**A** and **B**) Reconstructed cryo-ET tomographic slices of THP-1 cells, (**A**), and iBMDMs, (**B**), expressing WT ASC at different timepoints after stimulation with nigericin, or without stimulation (control). Scale bars, 200 nm. (**C**) Cryo-ET tomographic slices showing how the apex angles of mitochondrial cristae were measured for the quantitative analysis in **Fig. 4B**. Scale bars, 100 nm. (**D**) Cryo-ET tomographic slices showing how mitochondrial inter-cristae spacings were measured for the quantitative analysis in **Fig. 4B**. Scale bars, 100 nm.

**A**

| iBMDM type | Nigericin stimulation (min) | Gap diameter (nm) | Gap depth in z (nm) |
|---|---|---|---|
| ASC-mCerulean | 30 | 10.5 | 10.9 |
| ASC-mCerulean | 30 | 14.6 | 12.2 |
| ASC-mCerulean | 30 | 18.2 | 13.6 |
| ASC-mCerulean | 30 | 10.6 | 13.6 |
| WT | 60 | 20.4 | * |
| WT | 60 | 17.4 | 17.1 |

*gap near the end of Z stack

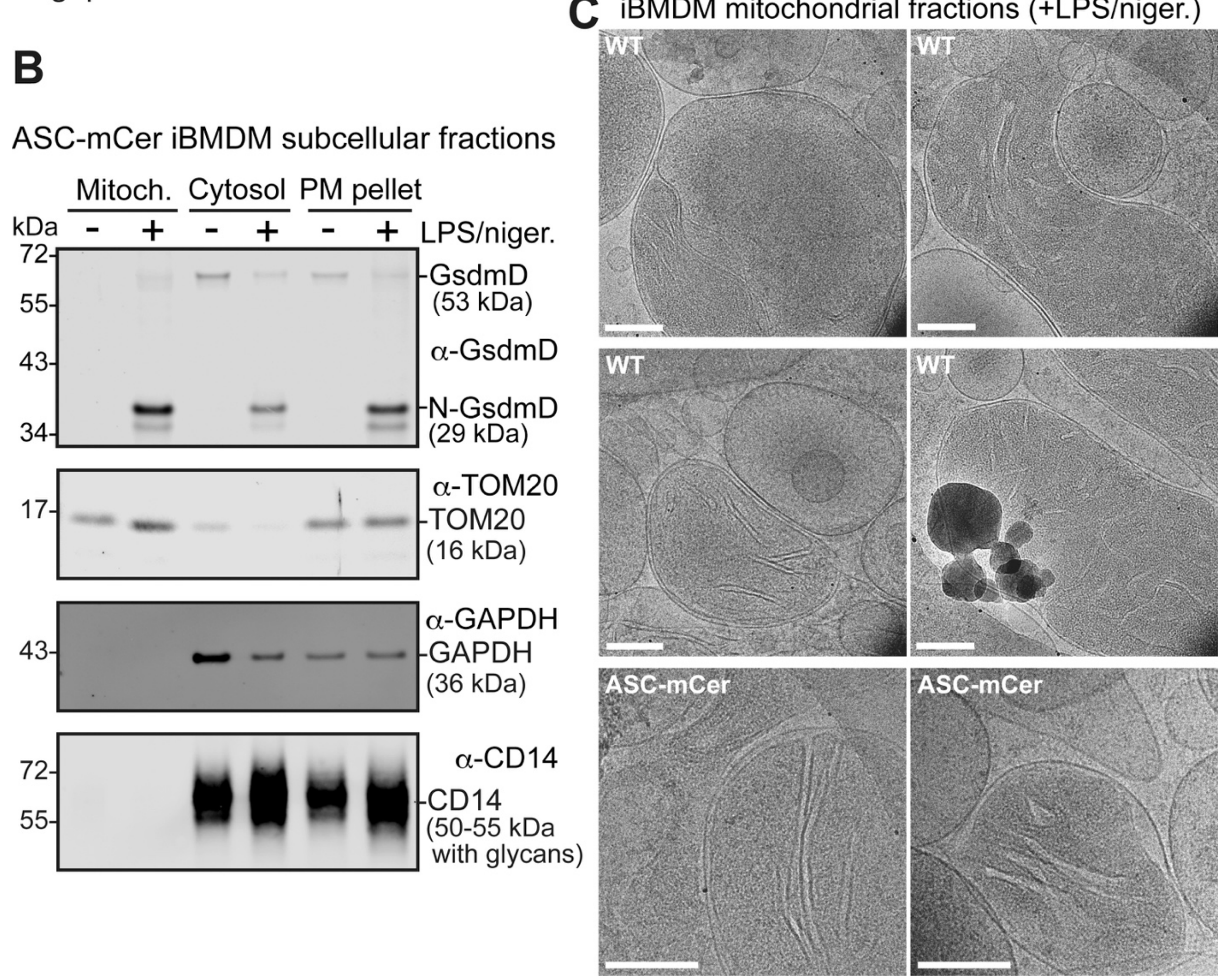


**Fig. S7. Mitochondrial outer-membrane pore sizes and purification of the mitochondrial subcellular fraction.** (**A**) Dimension of outer mitochondrial membrane pore in cryo-ET reconstructions of iBMDMs stimulated with LPG and nigericin. Two of the pores from ASC-mCerulean iBMDMs and both pores from WT iBMDMs are shown in **Fig. 5, A** and **B**. (**B**) Immunoblots of subcellular fractions of ASC-mCerulean iBMDMs 60 min after stimulation with LPS and nigericin. The GsdmD N-terminal domain (N-GsdmD) is enriched in the mitochondrial fraction. Shown below are immunoblots for Tom20 (a mitochondrial protein), GAPDH (a cytosolic protein), and CD14 (a plasma membrane protein). (**C**) Cryo-EM images of the purified mitochondrial fraction. Scale bars, 200 nm.

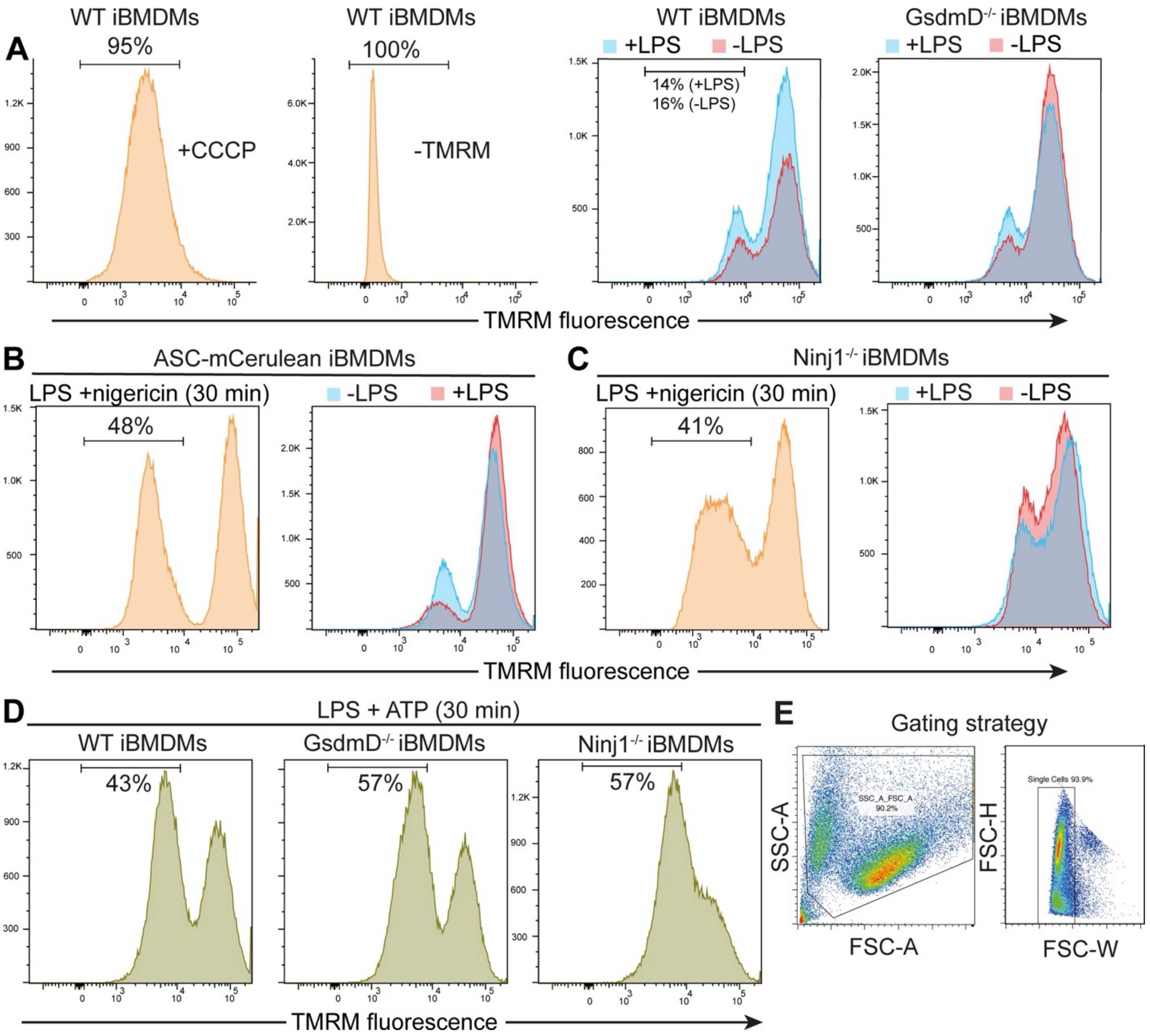


**Fig. S8. GsdmD contributes to mitochondrial membrane potential loss following NLRP3 activation.** (**A**) Flow cytometry controls supporting **Fig. 5, C** and **D**. From left to right: cells treated with carbonyl cyanide 3-chlorophenylhydrazone (CCCP), a mitochondrial uncoupler, and stained with TMRM (positive control); cells without TMRM staining (negative control); WT iBMDMs stained with TMRM with or without LPS priming (also shown in **Fig. 5C**); GsdmD$^{-/-}$ iBMDMs stained with TMRM with or without LPS priming. (**B**) 48% of ASC-mCerulean iBMDMs had loss of mitochondrial membrane potential following LPS priming and 30 min nigericin stimulation. Background controls without nigericin stimulation, with or without LPS priming are shown. (**C**) 41% of Ninj1$^{-/-}$ iBMDMs had loss of mitochondrial membrane potential following LPS priming and 30 min nigericin stimulation. Controls without nigericin stimulation, with or without LPS priming are shown. (**D**) Loss of mitochondrial membrane potential following priming with LPS and stimulation with ATP. (**E**) Gating strategy used to identify cells with loss of mitochondrial membrane potential. Side scatter area (SSC-A) and forward scatter area (FSC-A) were used to exclude debris. Forward scatter height (FSC-H) and forward scatter width (FSC-W) were used to exclude multiplet cells.

**Movie S1.**
Live-cell fluorescence imaging of iBMDMs stimulated with nigericin in the absence or presence of caspase-1 inhibitor Z-VAD-FMK (left and right, respectively).

**Movie S2.**
Z-stack series of reconstructed cryo-ET volume of the ASC-mCerulean punctum shown in **Figs. 1** and **2** overlaid with the 3-D segmented model shown in **Fig. 2**.

**Movie S3.**
Distribution of BODIPY TR ceramide during ASC speck formation in live iBMDMs expressing ASC-mCerulean (left), or costained with FAM-FLICA (right).

**Movie S4.**
Immunofluorescence micrograph Z-stack series of LPS-primed WT iBMDMs without nigericin stimulation (left), and after 30 min nigericin stimulation (right). Images from these Z-stack series are shown in **Fig. 3B**. Anti-NLRP3 partially colocalizes with anti-TGN38, and anti-TGN38 fluorescence disperses in the nigericin-stimulated cells. Pink, anti-NLRP3. Yellow, anti-TGN38. Scale bars, 10 μm.

**Data S1. (separate file)**
Source data for fluorescence recovery after photobleaching (FRAP; **Fig. 2G**).

**Data S2. (separate file)**
Source data for quantification of BODIPY TR ceramide fluorescence at sites of speck formation (examples shown in **Movie S3**).

**Data S3. (separate file)**
Source data for quantitative analysis of the morphological parameters of mitochondria in 3-D segmented surfaces of cryo-ET tomograms (**Fig. 4C**).

**Data S4. (separate file)**
Source data for **Fig. 5** and **Fig. S7**: quantitative analysis of the cryo-ET density at an outer mitochondrial membrane gap site (**Fig. 5B**); and uncropped Western blots (**Fig. 5E** and **Fig. S7B**).
